# Arabidopsis AGO1 N-terminal Poly-Q domain promotes phase separation and association with stress granules during heat stress

**DOI:** 10.1101/2023.10.12.562039

**Authors:** Aleksandar Blagojevic, Patricia Baldrich, Marlene Schiaffini, Esther Lechner, Nicolas Baumberger, Philippe Hammann, Taline Elmayan, Damien Garcia, Hervé Vaucheret, Blake C. Meyers, Pascal Genschik

**Affiliations:** Institut de Biologie Moléculaire des Plantes, CNRS, Université de Strasbourg, 12, rue du Général Zimmer, 67084 Strasbourg, France; Donald Danforth Plant Science Center, Saint Louis 63132, MO, USA; Plateforme Protéomique Strasbourg Esplanade du CNRS, Université de Strasbourg, 67084 Strasbourg, France; Université Paris-Saclay, INRAE, AgroParisTech, Institut Jean-Pierre Bourgin (IJPB), 78000 Versailles, France; Division of Plant Science and Technology, University of Missouri, Columbia, MO 65211, USA

## Abstract

In *Arabidopsis thaliana*, ARGONAUTE1 (AGO1) plays a central role in microRNA (miRNA) and small interfering RNA (siRNA)-mediated silencing. Nuclear AGO1 is loaded with miRNAs and exported to the cytosol where it associates to the rough ER to conduct miRNA-mediated translational repression, mRNA cleavage and biogenesis of phased siRNAs. These latter, as well as other cytosolic siRNAs, are loaded into cytosolic AGO1, but in which compartment this happens is not known. Moreover, the effect of stress on AGO1 localization is still unclear. Here, we show that a 37°C heat stress (HS) promotes AGO1 protein accumulation in cytosolic condensates where it co-localizes with components of siRNA bodies and of stress granules (SGs). AGO1 contains a prion-like domain in its poorly characterized N-terminal Poly-Q domain, which, is sufficient to undergo phase separation, independent of the presence or absence of SGS3. HS only moderately affects the small RNA repertoire, the loading of AGO1 by miRNAs and the signatures of target cleavage, suggesting that its localization in condensates protects AGO1 rather than promotes or impairs its activity in reprograming gene expressing during stress. Collectively, our work shed new light on the impact of high temperature on a main effector of RNA silencing in plants.

## INTRODUCTION

In eukaryotes, gene silencing is essential for development and plays important roles in response to stresses and for the epigenetic control of transposable elements (Ghildiyal and Zamore, 2009). At the molecular level, RNA silencing involves the processing of double-stranded (ds)RNA by Dicer-like RNase III enzymes, into small RNAs (sRNAs) ranging from 21 to 30 nucleotides (nt) in length (21 to 24-nt in plants). Small RNA duplexes are then incorporated into a protein complex called RISC (RNA-induced silencing complex); RISC always contains a member of the highly conserved Argonaute (AGO) protein family (Hutvagner and Simard, 2008)(Meister, 2013). In mammals, Ago2 is the only catalytic AGO protein, which upon loading with sRNAs, can engage in mRNA degradation and translational repression. Ago2 has been reported to localize to different membrane compartments, such as the Golgi body, the endoplasmic reticulum (ER) and multivesicular bodies, MVBs (Cikaluk et al., 1999) (Tahbaz et al., 2001) (Gibbings et al., 2009) (Lee et al., 2009). While the functional relevance of some of these subcellular localizations still requires elaboration, it is clear that miRNA- and siRNA-loaded Ago2 protein physically associates with the cytosolic side of the rough ER membrane to exert RNA silencing activity (Stalder et al., 2013). The association of Ago2 to MVBs has also been linked to silencing (Gibbings et al., 2009) (Lee et al., 2009), as this compartment seems required for RISC disassembly and may be involved in Ago2 secretion and/or lysosomal degradation. Besides these organelles, mammalian AGO proteins localize at specific non-membranous bodies, called P-bodies (Sen and Blau, 2005) (Liu et al., 2005a) (Liu et al., 2005b) (Pillai et al., 2005) (Jakymiw et al., 2005) (Jagannath and Wood, 2009)(Hubstenberger et al., 2017). However, P-bodies are not required for RNA silencing (Chu and Rana, 2006) (Eulalio et al., 2007). Under stress conditions Ago2 as well accumulates in other cytosolic bodies known as stress granules (SGs) (Leung et al., 2006). Understanding the subcellular localization, trafficking, and function of AGO proteins under stress represent important current challenges.

In the model plant *Arabidopsis thaliana* (hereafter Arabidopsis), AGO1 plays a central role in both siRNA- and miRNA-directed silencing (Fagard et al., 2000)(Vaucheret et al., 2004) (Mi et al., 2008). AGO1 loaded with sRNA mediates cleavage of target transcripts (Baumberger and Baulcombe, 2005) and is required for translational repression of at least a fraction of them (Brodersen et al., 2008)(Li et al., 2013). In line with its essential role in RNA silencing, *ago1* mutants or depletion of the AGO1 protein severely compromises plant development and affects cell division (Bohmert et al., 1998)(Morel et al., 2002)(Trolet et al., 2019). In addition, AGO1 is an important regulator of antiviral defense, as its mutation enhances susceptibility to different RNA viruses (Morel et al., 2002)(Azevedo et al., 2010)(Clavel et al., 2021).

At the subcellular level, it was recently reported that AGO1 carries both nuclear localization (NLS) and nuclear export signal (NES) signals. Unloaded cytosolic AGO1 has exposed NLS and hidden NES, and is imported in the nucleus where it can be loaded with miRNAs, thus exposing its NES, which promotes its export to the cytosol in the form of miRNA- AGO1 complexes (Bologna et al., 2018). In the nucleus, AGO1 has also been linked to other functions such as transcription and RNA-mediated DNA repair (reviewed in (Bajczyk et al., 2019)). Once in the cytosol, AGO1 appears in membrane-free (soluble) and membrane-bound forms, the latter being mainly the ER (Brodersen et al., 2012)(Li et al., 2013)(Li et al., 2016)(Michaeli et al., 2019). In fact, AGO1 association to the rough ER is the site of miRNA-mediated translational repression (Li et al., 2013) and biogenesis of phased siRNA on membrane-bound polysomes (Li et al., 2016). Cytosolic AGO1 is also loaded with siRNAs produced in the cytosol, and the resulting complexes execute mRNA cleavage, which is exemplified in antiviral defense and posttranscriptional gene silencing (PTGS) (Carbonell and Carrington, 2015). Of note, PTGS also involves the production of secondary siRNAs via the action of the RNA DEPENDENT PROTEIN RNA POLYMERASE 6 (RDR6) and SUPPRESSOR OF GENE SILENCING 3 (SGS3), two components that reside in cytosolic foci called siRNA-bodies (Jouannet et al., 2012). Whether Arabidopsis AGO1 associates with siRNA-bodies or other cytoplasmic cellular compartments requires more investigation. Moreover, its subcellular distribution when plants encounter stress remains unclear.

Temperature is an important environmental factor that affects plant growth and development and heat stress (HS) causes significant reduction in crop yield and quality (Battisti and Naylor, 2009)(Ohama et al., 2017). Previous work has shown that growing Arabidopsis plants permanently at 30°C inhibits PTGS mediated by the RDR6/SGS3 siRNA pathway (Zhong et al., 2013). However, a recent report suggested that a short heat stress (HS) at 42°C stimulated the formation of siRNA bodies via SGS3 phase separation, leading to a massive production of endogenous siRNAs (Tan et al., 2023). In the present study we questioned how HS and its recovery affect AGO1 subcellular distribution and activity in RNA silencing. To avoid artifacts resulting from overexpression and/or transient expression assays, we performed most of our studies with Arabidopsis plants carrying genomic constructs of AGO1 and of other cellular compartment markers fused to fluorescent proteins and expressed under the control of their own regulatory elements. Using Confocal Laser Scanning Microscopy (CLSM) imaging of root meristematic cells, we show that upon HS at 37°C, AGO1 dynamically associates in condensates corresponding to stress granules (SGs). Besides SG proteins, the interactome of AGO1 upon HS also revealed P-body and RNA decay components. The recruitment of AGO1 in these condensates does not require SGS3; rather we show that the N-terminal Poly-Q domain of AGO1 has the ability to undergo phase separation, both *in vitro* and *in vivo*. Using transcriptomics, we observed that while protein-coding genes are promptly responding to the HS and recovery condition, sRNAs are not impacted by short-term HS treatment. Additionally, we observed that miRNA accumulation, AGO1 loading or cleavage are not substantially affected by HS, suggesting that these AGO1 functions are not linked to AGO1 localization in stress granules.

## RESULTS

### AGO1 protein accumulates and dynamically associates with siRNA bodies during HS

Under normal growth conditions and as previously reported, a functional GFP-AGO1 fusion protein is mainly localized in the cytoplasm of Arabidopsis root tip cells (Fig. 1A) (Derrien et al., 2012)(Michaeli et al., 2019)(Bologna et al., 2018). However, upon 30 min HS treatment at 37°C, AGO1 subcellular distribution substantially changed, with the appearance of GFP-AGO1 cytosolic foci that persisted for at least for 4 hours during continuous HS (Fig. 1A). Nonetheless, after HS, most of these foci disappeared during 2 hours of temperature recovery at 22°C, and the diffuse AGO1 cytosolic pattern was restored (Fig. 1B), indicating that HS-dependent AGO1 cellular relocalization is a dynamic process. We next investigated if HS affects AGO1 expression level and observed that the amount of AGO1 protein increased during HS in a transient manner (Fig. 1C). The HS-dependent AGO1 protein accumulation did not correlate with an increase of AGO1 steady-state mRNA level, but correlated with a lower accumulation of miR168, which guides AGO1 mRNA cleavage and translation repression (Vaucheret et al., 2004)(Mallory et al., 2009) (Várallyay et al., 2010) (Fig. 1C). Thus, heat-induced AGO1 protein accumulation occurs at the post-transcriptional level.

**Figure 1:**
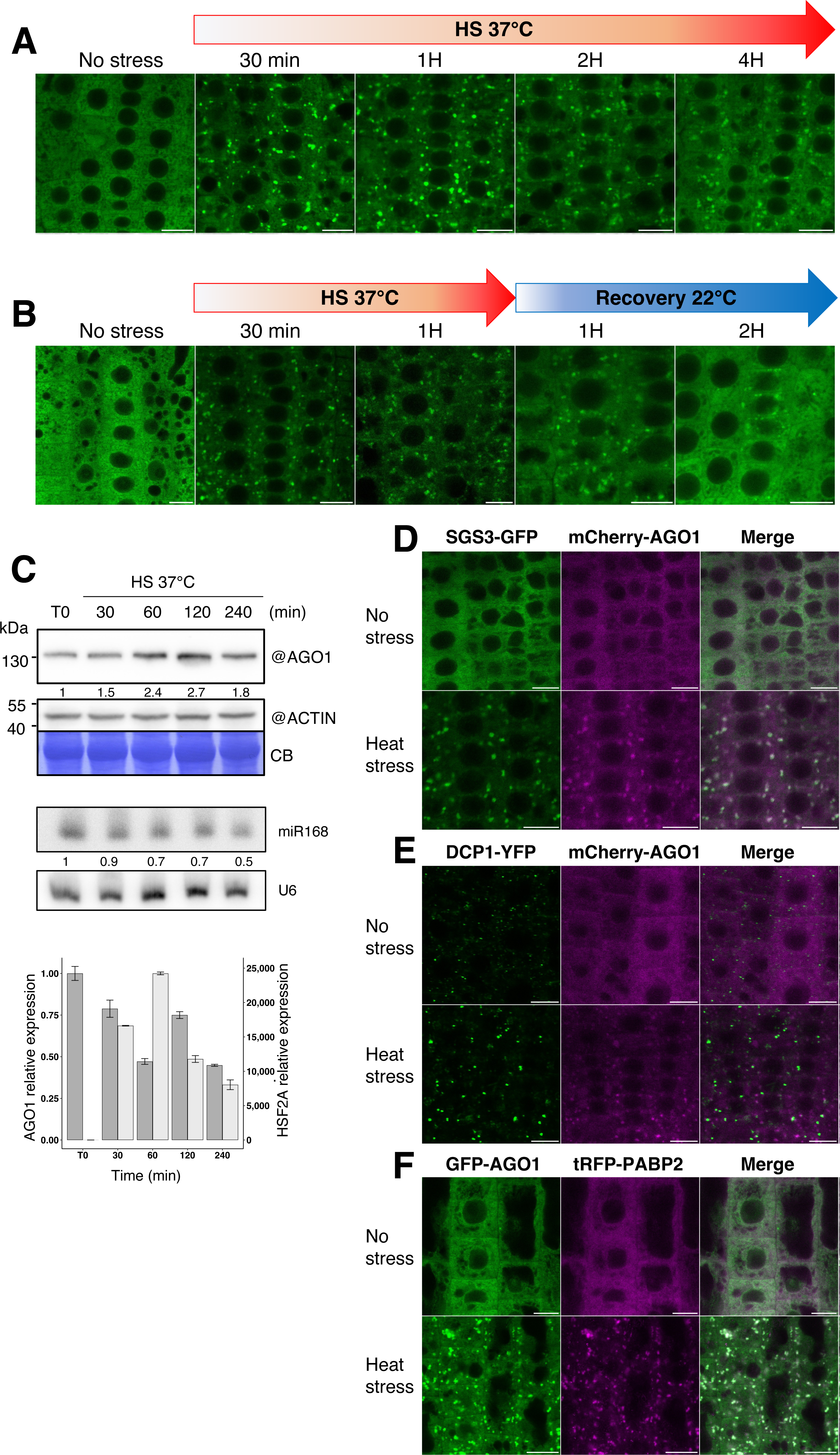
AGO1 protein accumulation and colocalization with the stress granules under HS. **(A-B)** Dynamics of accumulation and disappearance of AGO1 foci during HS. CLSM imaging of 7 day old Arabidopsis root tip cells of the pAGO1:GFP-AGO1 transgenic line subjected to continuous HS (A) or 1 hr HS at 37°C followed by 2 hrs recovery at 22°C (B). Bar = 10 µm. Objective 40X, oil immersion. **(C)** AGO1 protein and RNA accumulation during HS. Kinetic analysis of 7 day old Arabidopsis Col-0 seedlings grown on MS medium and subjected to HS. Upper panel: AGO1 protein was detected by western blot with a specific antibody. Coomassie blue (CB) staining and ACTIN were used as loading controls. AGO1 signal was quantified by ImageJ and normalized to the corresponding ACTIN signal. Numbers below the AGO1 panel are indicated as relative to non-stressed Col-0 (T0) set at 1.0. Middle panel: sRNA gel blot analysis of the steady-state accumulation of the indicated miRNA taken from the same material as above. U6 RNA level was used as a loading control. miRNAs signals were quantified by ImageJ and normalized to the corresponding U6 signal. Numbers below panels are indicated as relative to non-stressed Col-0 (T0) set at 1.0. Bottom panel: RT-qPCR analysis of AGO1 and HSFA2 transcript levels. RNA samples were extracted from the same material as above. Bars indicate the mean expression of three technical replicates, and error bars indicate the SD. **(D)** AGO1 colocalizes with SGS3 in foci under heat-stressed conditions. CLSM imaging of 5 day old Arabidopsis pAGO1:mCherry-AGO1 x pRDR6:SGS3-GFP root tip cells before and after 37°C HS of 30 minutes. Experiments were performed in triplicate with three roots per replicate. Bar = 10 µm. Objective 40X, oil immersion. Foci number after HS for SGS3-GFP = 842 ± 446; PCC= 0.77 ± 0.06. Foci number after HS for mCherry-AGO1 = 789 ± 314; PCC= 0.70 ± 0.10. **(E)** DCP1 does not colocalize with AGO1 under heat stressed conditions. CLSM imaging of 5 day old pDCP1:DCP1-YFP *dcp1-3* x pAGO1:mCherry-AGO1 Arabidopsis root tip cells before and after a 37°C HS of 30 minutes. Experiments were performed in triplicate with three roots per replicate. Bar = 10 µm. Objective 40X, oil immersion. Foci number before HS for DCP1-YFP = 743 ± 345; PCC= 0.7 ± 0.10. Foci number after HS for DCP1-YFP = 690 ± 241; PCC= 0.31 ± 0.12. Foci number after HS for mCherry-AGO1 = 703 ± 209; PCC= 0.32 ± 0.06. **(F)** AGO1 colocalizes with PABP2 in foci under heat stressed conditions. CLSM imaging of 5 day old Arabidopsis pAGO1:GFP-AGO1g *ago1-27* x pPABP2:tRFP-PABP2 root tip cells before and after 37°C HS of 30 minutes. Experiment performed in triplicate with three roots per replicate. Bar = 10 µm. Objective 40X, oil immersion. Foci number after HS for GFP-AGO1 = 1777 ± 424; PCC= 0.76 ± 0.11. Foci number after HS for tRFP-PABP2 = 2257 ± 581; PCC= 0.77 ± 0.09.

To identify intracellular organelles with which AGO1 may associate during HS, we crossed AGO1 reporter lines with Arabidopsis lines expressing different cytoplasmic compartment markers (Golgi, the trans-Golgi network/early endosome (TGN/EE) and multivesicular body (MVB), allowing dual subcellular colocalization studies (see Table S1). Of particular interest are MVBs as AGO1 was recently found in extracellular vesicles, originating from this compartment (He et al., 2021). Moreover, MVBs also play important roles in trafficking proteins to the vacuoles (Cui et al., 2016) and vacuolar degradation of AGO1, at least in a viral context, has previously been reported in plants (Derrien et al., 2012)(Michaeli et al., 2019). Nonetheless, the fluorescence intensity between the AGO1 and MVB channels did not correlate in Arabidopsis root tip cells, confirmed by the low value of the Pearson coefficients (PPCs) (Fig. S1A), indicating that GFP-AGO1 does not appear to colocalize with MVBs. Similarly, and despite a clear reorganization of these membranous organelles during HS, no colocalization of AGO1 with the TGN/EE and Golgi was observed (Fig. S1B-C).

Next, we examined whether AGO1 associates with siRNA bodies upon HS. siRNA bodies are non-membranous cytoplasmic structures characterized by the presence of the proteins SGS3 and RDR6 (Elmayan et al., 2009) (Kumakura et al., 2009). AGO7, which participates in the production of trans-acting siRNA (ta-siRNA), is also found in siRNA bodies (Jouannet et al., 2012). Notably siRNA body formation was previously shown to be distinct from P-bodies and to dramatically increase under HS in leaves of *Nicotiana tabacum* (Jouannet et al., 2012) (Martínez de Alba et al., 2015). Thus, we crossed the pAGO1:mCherry-AGO1 line with the pRDR6:SGS3-GFP line (Table S1). Confocal imaging of Arabidopsis root cells indicated that under normal growth conditions the SGS3-GFP signal was mainly diffuse and that only a few siRNA bodies were present (Fig. 1D). Once we transferred seedlings for 30 min at 37°C, we observed a massive accumulation of foci, in which mCherry-AGO1 and SGS3- GFP were colocalized, as supported by the high value of PPCs for both proteins (Fig. 1D; Movie S1).

### HS-induced AGO1 bodies colocalize with stress granules

We next investigated the cellular interactions between AGO1 containing foci and two other membrane-less ribonucleoparticles (RNPs) composed of an association of mRNA and proteins: P-bodies and stress granules (SGs). P-bodies are ubiquitously present in cells and contain RNA decapping enzymes and exonucleases and are known to play important roles in mRNA catabolism (Chantarachot and Bailey-Serres, 2018). Contrary to P-bodies, stress granules (SGs) are not present in non-stressed cells and form from mRNAs that become stalled in translation initiation during environmental stress such as HS (Riggs et al., 2020)(Chantarachot and Bailey-Serres, 2018). We first visualized P-bodies and SGs in our experimental conditions, by using an Arabidopsis transgenic line expressing DCP1-YFP and tRFP-PABP2, respectively, and confirmed that both entities do not colocalize under HS as previously reported (Fig. S2A) (Jouannet et al., 2012) (Martínez de Alba et al., 2015). Note that the mobility of P-bodies changes with stress; while they are very mobile under normal growth conditions, they become more static under HS (Movies S2 and S3) and are often found in close proximity to SGs, potentially allowing for cross-talk of the two entities (Martínez de Alba et al., 2015)(Kumakura et al., 2009).

We next investigated the localization of mCherry-AGO1 with respect to the DCP1-YFP maker of P-bodies (Fig. 1E). In the absence of stress, we did not observe a clear co-localization of the proteins, suggesting that AGO1, at least in Arabidopsis root tip cells, is not a major component of P-bodies. Moreover, during HS, the two proteins localized to distinct, not overlapping foci. On the other hand, GFP-AGO1 fully colocalized with the SG marker tRFP- PABP2 under HS Fig. 1F). This was also observed for SGS3-GFP (Fig. S2B) and is in line with previous reports showing colocalization of siRNA bodies and SG markers after HS (Jouannet et al., 2012)(Tan et al., 2023). Thus, the size of HS-induced AGO1-containing bodies, their low mobility and localization with PAB2 suggest that AGO1 is compartmentalized in SG condensates during HS.

Notably, SGs are sensitive to drugs inhibiting translation elongation such as cycloheximide (CHX), which prevents their formation in mammalian cells after drug treatment during stress (Kedersha et al., 2000)(Ivanov et al., 2019). Thus, we investigated whether this is also the case for the HS-induced AGO1-containing bodies. Five-day old seedlings expressing both GFP-AGO1 and tRFP-PABP2 were subjected to 37°C HS for 30 minutes in the absence or presence of 100µM CHX (Fig. 2). As expected, tRFP-PABP2 shows a diffuse cytosolic distribution in the presence of CHX, indicating the disassembly of SGs. Remarkably, HS-induced AGO1-containing bodies were still visible in the presence of CHX, though of smaller size (Fig. 2). Similar results were also observed for an Arabidopsis line expressing both SGS3-GFP and tRFP-PABP2 (Fig S3), for which SGS3 decorated bodies were still visible under HS in the presence of CHX, in line with the recent report of (Tan et al., 2023). Thus, while components of siRNA bodies and SG perfectly co-localize under HS, they also show some distinct features.

**Figure 2:**
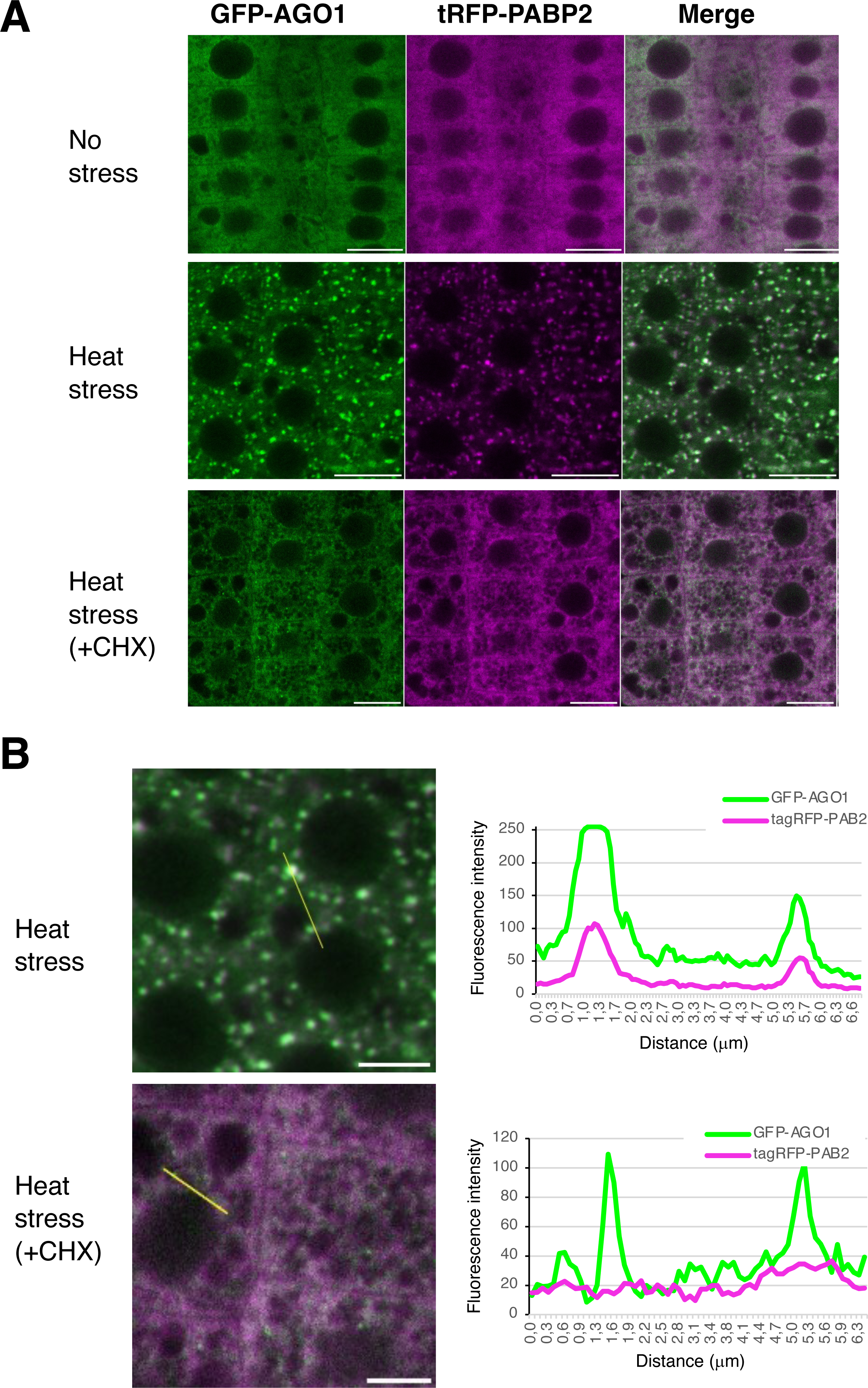
Cycloheximide inhibits stress granule formation, but allows the persistence of small AGO1 foci. **(A)** CLSM imaging of 5 day old Arabidopsis pAGO1:GFP-AGO1g *ago1-27* x pPABP2:tRFP- PABP2 root tip cells before HS (upper panels), after 37°C HS of 30 minutes (middle panels) and after 37°C HS of 30 minutes in the presence of 100µM CHX). Experiment performed in duplicate with three roots per replicate. Bar = 10 µm Objective 40X, oil immersion. Colocalization of GFP-AGO1 and tRFP-PABP2 after HS in CHX-untreated cells: PCC = 0.96 Colocalization of GFP-AGO1 and tRFP-PABP2 after HS in CHX treated cells: PCC = 0.52 **(B)** Signal intensity distribution of the total amount of pixels at the x axis shown in CHX- untreated and CHX-treated cells after 30 minutes of HS at 37°C shown in (A).

### Besides SG proteins some P-bodies and RNA decay components are also part of the HS-induced AGO1 interactome

To identify the interaction network of AGO1 during HS, we immunoprecipitated GFP-AGO1 protein from Arabidopsis pAGO1:GFP-AGO1 *ago1-27* seedlings after 1 hour at 37°C and performed mass spectrometry analysis. Proteins significantly enriched in the GFP-AGO1 IP were highlighted by a statistical analysis, calculating normalized fold changes and adjusted p values (Fig. 3A and Table S2). In accordance with our microscopic observations, a large category of proteins predominantly enriched in the immunoprecipitation (IP) corresponded to SG components, including PAB2, RNA-binding protein 47A-C (RBP47A-C), RNA helicases (RH6/8/11/52/53), G3BP1, and TSN1/2, among others. In total, 72 proteins significantly enriched in our HS-induced AGO1 IP were shared with the reported Arabidopsis SG proteome (Kosmacz et al., 2019).

**Figure 3:**
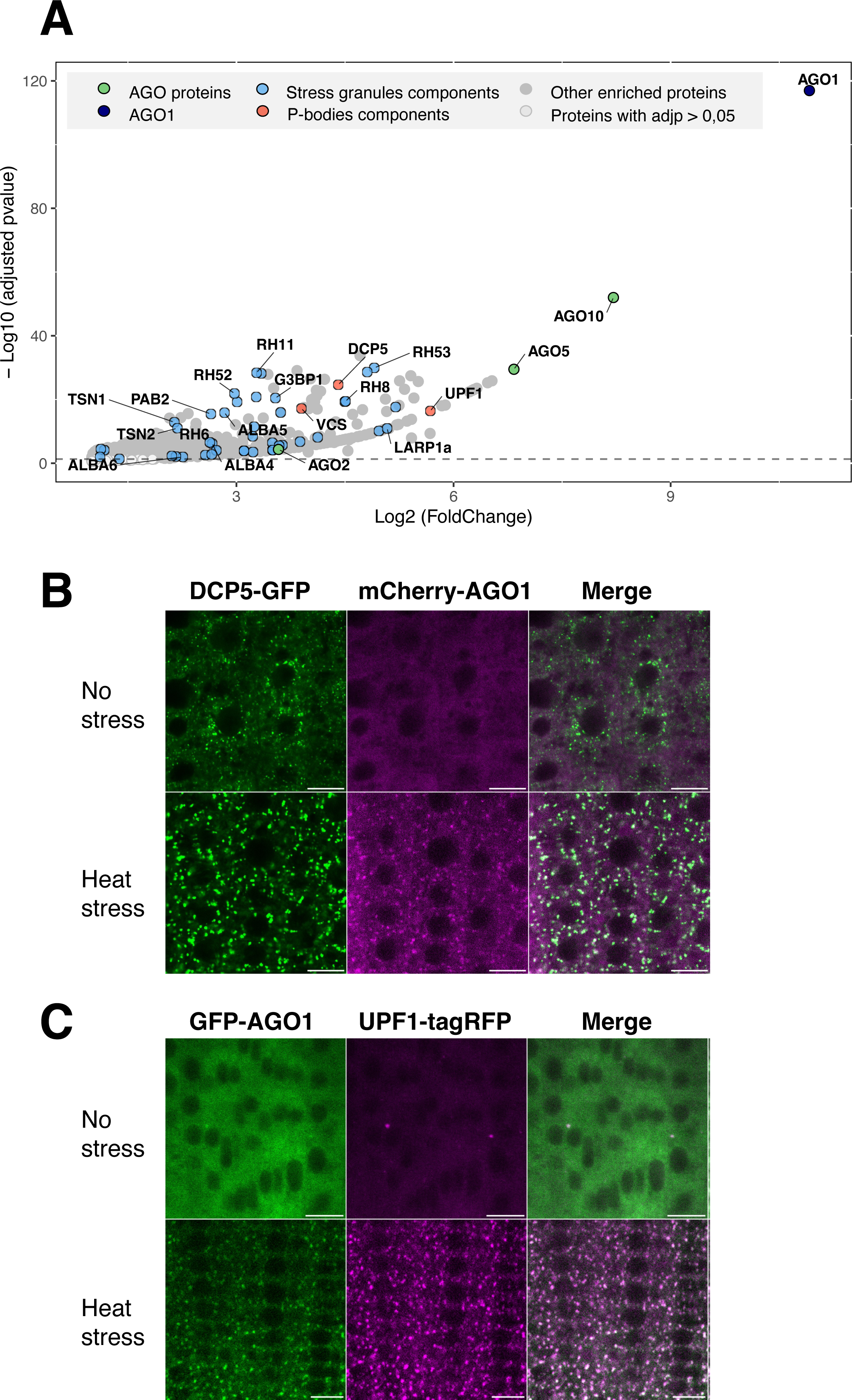
AGO1 interactome and its colocalization with P-body and RNA decay components under heat stress. (A) AGO1 interactome revealed by immunoprecipitation and mass spectrometry. 7 day old Arabidopsis pAGO1:GFP-AGO1g *ago1-27* and Col-0 seedlings were subjected to 1 hr HS at 37°C. We compared 12 samples (pAGO1:GFP-AGO1g *ago1-27*) from 3 independent biological replicates to 12 control samples (Col-0 seedlings subjected to 1 hr HS at 37°C). Volcano plot shows the enrichment of proteins co-purified with the GFP-AGO1 bait compared with Col-0 controls. The y and x axes display log2 values from adjusted p values and fold changes, respectively. The horizontal dashed line indicates the threshold above which proteins are significantly enriched (adjusted p values < 0.05). Only AGO1-enriched proteins (log2FC > 1) are shown. Three color-coded functional clusters are highlighted. Enriched proteins are dark grey, cytoplasmic RNA granules related proteins are highlighted in blue, AGO proteins are in green, selected P-bodies markers are in red. The source data are available in Table S2. (B) DCP5 colocalize with AGO1 under heat stressed conditions. CLSM imaging of 5 day old Arabidopsis pDCP5:DCP5-GFP *dcp5-1* x pAGO1:mCherry-AGO1 root tip cells before and after 37°C HS of 30 minutes. Experiments were performed in triplicate with three roots per replicate. Bar = 10 µm. Objective 40X, oil immersion. Foci number after HS for DCP5-GFP = 544 ± 180; PCC= 0.58 ± 0.08. Foci number after HS for mCherry-AGO1 = 429 ± 198; PCC= 0.53 ± 0.09. (C) AGO1 colocalizes with UPF1 in both normal and heat-stressed conditions. CLSM imaging of 5 day old Arabidopsis pAGO1:GFP-AGO1 *ago1-27* x pUPF1:UPF1-tagRFP *upf1-5* root tip cells before and after 37°C HS of 30 minutes. Experiments were performed in duplicate with three roots per replicate. Bar = 10 µm. Objective 40X, oil immersion. Foci number before HS for UPF1-tagRFP = 69 ± 84; PCC= 0.67 ± 0.14. Foci number after HS for UPF1-tagRFP = 1344 ± 629; PCC= 0.76 ± 0.12. Foci number after HS for GFP-AGO1 = 1074 ± 658; PCC= 0.74 ± 0.13.

Surprisingly, we also identified some P-body and RNA decay components such as DCP5, VARICOSE, LARP1a and UP-FRAMESHIFT1 (UPF1) in the AGO1 interactome. As AGO1 does not colocalize with DCP1 (Fig. 1E), this raises the possibility that some, but not necessarily all components of P-bodies, might associate with HS-induced AGO1 condensates. To address this question, we generated Arabidopsis transgenic lines expressing mCherry- AGO1 in the pDCP5:DCP5-GFP *dcp5-1* background or GFP-AGO1 in pUPF1:UPF1-tagRFP *upf1-5* background (Table S1). Under non-stress conditions we did not observe co-localization between DCP5-GFP and mCherry-AGO1 (Fig. 3B) but the proteins did colocalize during HS. To confirm that DCP5 colocalizes with SG we also generated an Arabidopsis line expressing both DCP5-GFP and tRFP-PABP2. Under normal growing conditions only DCP5-GFP forms foci, likely corresponding to P-bodies, but under HS, DCP5-GFP co-localized with tRFP- PABP2 in SGs (Fig. S4A). For UPF1-tagRFP, we observed colocalization with GFP-AGO1 in few foci in the absence of HS, while under HS, both proteins wholly colocalized (Fig. 3C). Finally, we investigated the localization of SGS3-GFP with respect to UPF1-tagRFP and confirmed that during HS, both proteins partially colocalize (Fig. S4B), which is in accordance with previous observations showing partial colocalization of both proteins when transiently expressed in *Nicotiana benthamiana* (hereafter *N. benthamiana*) leaves (Moreno et al., 2013). Overall, these data indicate that HS promotes the association of AGO1 with SG condensates together with siRNA-bodies components and other proteins involved in RNA turnover, including the decapping-associated factor DCP5 and UPF1, a key component of nonsense-mediated mRNA decay (NMD).

### A short HS at 37°C has limited effect on sRNA accumulation and their loading into AGO1

Next, we investigated how AGO1 loading and activity is affected by HS. To address this question, 7-day old Col-0 seedlings grown at 22°C were subjected to HS at 37°C for 1 hr, followed by 2 hrs recovery at 22°C. We performed sequencing analyses on coding RNAs, and total and AGO1-associated sRNAs. The transcriptomic analysis of the same RNA from wild-type Col-0 plants exposed to heat stress and recovery revealed that a total of 9724 genes were differentially expressed in at least one of our three comparisons (Fig. S5A; UpsetPlot). Out of this subset of genes, only 735 (or 7.5%, red) appear to be differentially expressed in all comparisons. The most transcriptional changes happened during the recovery phase since 6626 (or ∼68%) genes appeared to be differentially expressed when comparing recovery to control, recovery to HS, or common to both comparisons (Fig. S5A, blue). These genes were enriched in the “Response to Stress” GO category. As a reference subset, out of the 68 loci identified in TAIR (version 10) as heat hock proteins (HSPs) and heat shock factors (HSFs), 42 were differentially expressed in response to heat, including nuclear, mitochondrial and chloroplastic HSPs (Fig. 4A). Most of these heat shock related genes were highly expressed when comparing HS versus control plants and remained at high levels when comparing recovery to control plants. However, when comparing recovery to HS plants, we observed their downregulation in accordance with the exit of the transcriptional HS response during recovery.

**Figure 4:**
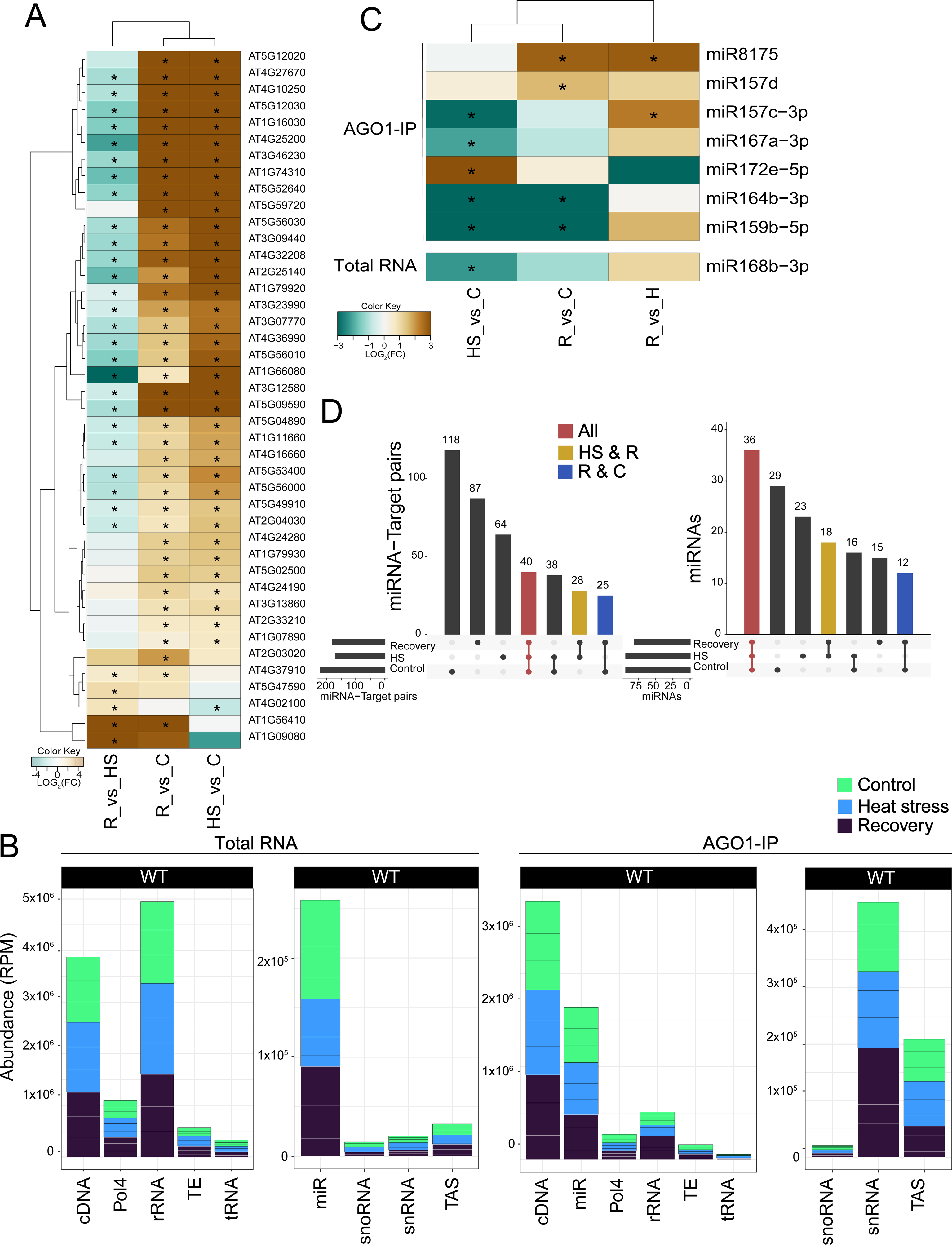
sRNA accumulation and loading into AGO1 during HS and recovery. (A) Heat shock-related genes are differentially expressed in response to treatments. Heat map of differentially expressed genes encoding heat shock related proteins annotated in the TAIR10 genome. Brown indicates enrichment and teal indicates depletion in each of the three comparisons. C, Control; HS, Heat Stress; R, Recovery. (B) sRNAs are not affected by HS. The sRNAs that mapped to the genome were categorized by origin annotated in TAIR10 and plotted by relative abundance in RPM. tRNA, Transfer RNAs; cDNA, complementary DNA; Pol IV, products dependent on RNA polymerase IV; snRNAs, small nuclear RNAs; snoRNA, small nucleolar RNA; TAS, trans-acting siRNA; rRNA, ribosomal RNAs. (C) Eight miRNAs differentially accumulated in response to heat treatment and recovery. Heat map of differentially expressed miRNAs in AGO1-IP (upper panel) and total RNA (lower panel). Brown indicates enrichment and teal indicates depletion in each of the three comparisons. C, Control; HS, Heat Stress; R, Recovery. (D) Upset plot representing the miRNA-target pairs (left panel) and miRNAs only (right panel) identified in nanoPARE sequencing for all three comparisons. For each upset plot, the bottom left shows the number of miRNA target pairs in each comparison as a horizontal histogram, the bottom right shows the intersection matrix and the upper right shows the size of each combination as a vertical histogram. The red color marks common pairs to all three comparisons, the yellow color marks common pairs between HS and recovery, and the blue lines mark common pairs between recovery and control.

To assess the impact of HS on sRNAs, we generated sRNA libraries from total RNA and AGO1-IP on the same biological replicates used for the transcriptomic analysis. The sRNA libraries had the expected size distributions with peaks at 21- 23- and 24-nt for total RNA and 20- and 21-nt for AGO1-IP, independent of the heat treatment (Fig. S5B). sRNAs that mapped to the genome were categorized by origin and plotted by relative abundance (Fig. 4B). Overall, no major differences in sRNA origin between treatments were observed. Moreover, there was no significant increase in any of the tasiRNA population (Fig. S5C). We also analyzed the differential accumulation (DA) of each miRNA in each sample. For total RNA, only one miRNA (miR168b-3p) exhibited DA when comparing HS to control (Fig. 4C). When analyzing the miRNAs loaded into AGO1, we found 7 miRNAs with DA in several samples. The majority of these miRNAs were downregulated during HS compared to control. We therefore conclude that neither the miRNA accumulation nor the loading in AGO1 changes dramatically in response to HS.

We next asked whether some miRNAs could target specific transcripts during HS and recovery. Thus, we generated nano-parallel analysis of RNA end (nanoPARE) libraries from the same material used for the sRNA-seq libraries. Considering only the miRNA-target PARE present in two out of three biological replicates, we found that most of the miRNA target signatures detected are unique to each group (control, HS, and recovery) (Fig. 4D, left panel). However, when comparing only miRNAs or targets that had a signature in nanoPARE, without considering their counterparts, we found that the majority of miRNAs were shared between all treatments (Fig. 4D, right panel), while the majority of targets were unique to each treatment (Fig. S5D). We conclude that there are few changes happening at the miRNA level and that the changes we observed might be happening at the transcript target level.

### AGO1 undergoes LLPS though its N-terminal domain

An intriguing question remains how does AGO1 accumulate in condensates under HS? Recent work revealed that SGS3 has two prion-like domains (PrLDs) that mediate liquid–liquid phase separation (LLPS) in siRNA body formation (Kim et al., 2021)(Tan et al., 2023). Interestingly, PrLDs have been predicted as protein motifs for Arabidopsis AGO1 and AGO2 (Chakrabortee et al., 2016), but not characterized yet. For AGO1, the unique PrLD is located in the N-terminal part of the protein in a region called the Poly-Q domain due to its high glutamine content (Fig. 5A). In fact, the Poly-Q domain encompasses an intrinsically disordered regions (IDR), which is predicted with good probability to undergo LLPS. To investigate the importance of the AGO1 N-terminal region for its subcellular localization, we engineered an AGO1 construct lacking the PrLD (AGO1(DPrLD)). The native version of AGO1 and the mutated variant were expressed as GFP fusion proteins in *N. benthamiana* leaves subjected, or not, to 30 min of HS at 37°C. In contrast to the stable expression of GFP-AGO1 in Arabidopsis, a significant number of foci could already be observed with the AGO1 (WT) construct even in the absence of HS in *N. benthamiana* (Fig. 5B). This is likely due to the stress resulting from the transient expression assay and/or the overexpression of AGO1. Even so, HS stimulated the formation of foci for both AGO1 constructs, indicating that this response does not depend on the sole PrLD.

**Figure 5:**
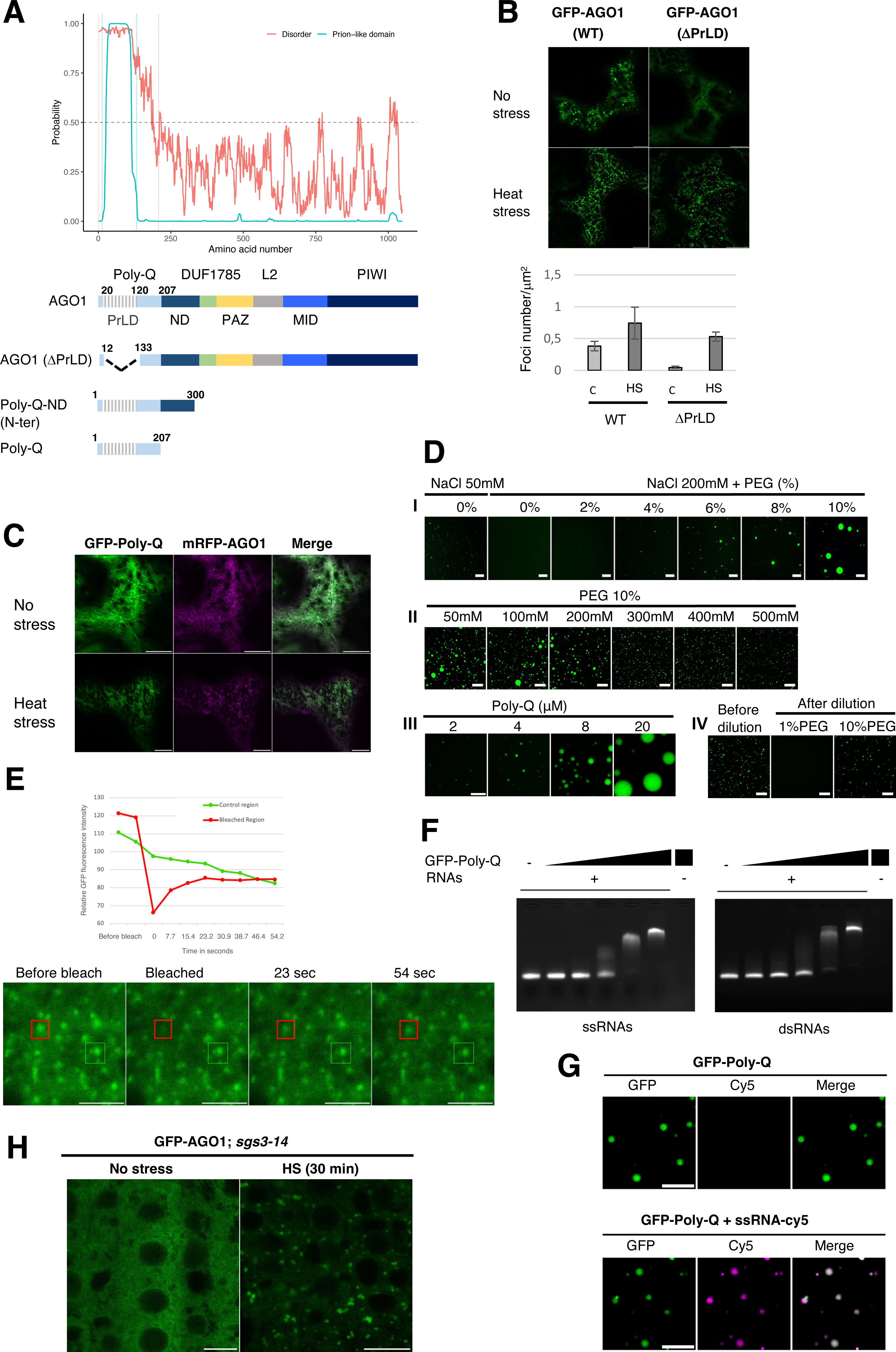
The N-terminal domain of AGO1 mediate HS-dependent LLPS. (A) Schematic representation of the AGO1 deletion constructs used to assay LLPS in *N. benthamiana*. Each construct contains a GFP at the N-terminus and is under the control of the CaMV 35S promoter. The name of each AGO1 protein domain is indicated. The Poly-Q domain at the N-terminus is predicted to be both prion-like and intrinsically disordered. Prion-like domain and intrinsically disordered region were predicted using respectively http://plaac.wi.mit.edu/, https://iupred2a.elte.hu/. (B) The deletion of AGO1 PrLD does not abolish the efficiency for LLPS. Representative CLSM imaging of *N. benthamiana* leaf epidermal cells transiently expressing GFP-AGO1 (AGO1) and AGO1(ΔPrLD) proteins without or with HS (30 minutes at 37°C). Bar = 10 µm. Objective 40X, oil immersion. Bottom panel, quantification of GFP-labelled bodies for each construct per cytosolic area unit (foci number/μm²). (C) The GFP-Poly-Q fusion protein colocalizes in the same foci as full-length mRFP-AGO1 under HS. CLSM imaging of *N. benthamiana* leaf epidermal cells transiently coexpressing p35S:GFP-Poly-Q and p35S:mRFP-AGO1 without or with HS (30 minutes at 37°C). Bar = 10 µm. Objective 40X, oil immersion. (D) AGO1 Poly-Q domain alone forms droplet like structures *in vitro.* (I) *In vitro* analysis of droplet formation by recombinant GFP-Poly-Q AGO1 protein at different PEG8000 concentrations in Hepes 25mM buffer pH7.5. Protein concentration 4μM. Scale bars, 20μm. Objective 40X. (II) *In vitro* analysis of droplet formation by recombinant GFP-Poly-Q AGO1 protein at different NaCl concentration in Hepes 25mM buffer pH7.5, PEG 8000 10%. Protein concentration 4μM. Scale bars, 50μm. Objective 20X. (III) *In vitro* analysis of droplet formation by recombinant GFP- Poly-Q AGO1 protein at different protein concentrations in Hepes 25mM buffer pH7.5, NaCl 200mM, PEG8000 10%. Scale bars, 20μm. Objective 40X. (IV) Reversion assay of droplets by 10-fold dilution of a concentrated (18μM) GFP-Poly-Q AGO1 in 25mM HEPES pH7.5, 200mM NaCl, 10% PEG 8000 in solutions with or without PEG. Scale bars, 50μm. Objective 20X. (E) FRAP assay of pAGO1:GFP-AGO1 ago1-27 Arabidopsis root tip cells subjected to HS (30 minutes at 37°C). The fluorescence intensity during the time course of recovery after photobleaching and microscopy images are indicated. Time 0 indicates the time of the photobleaching pulse. Red and green scares indicate the photobleached and control condensates respectively. Bar = 10 µm. Objective 40X, oil immersion. (F) Agarose-gel EMSA assay showing that GFP-Poly-Q domain binds ssRNAS and dsRNAs *in vitro*. RNAs are at 0,9μM, GFP-Poly-Q AGO1 protein ranges from 0,56 to 9μM by two-fold increment. (G) Colocalization of GFP-Poly-Q AGO1 protein with Cy5 labelled-ssRNA upon droplet formation in Hepes 25mM buffer pH7.5, NaCl 200mM, PEG8000 10%. Protein and RNAs are at 4μM. Scale bars, 10μm. Objective 40X. (H) AGO1 does not rely on SGS3 to form condensate during HS. CLSM imaging of 5 day old Arabidopsis root tip cells of the pAGO1:GFP-AGO1 x *sgs3-14* transgenic line subjected to 30 min HS at 37°C. Bar = 10 µm. Objective 40X, oil immersion.

Next, we investigated whether the AGO1 N-terminal region alone is sufficient to induce phase separation. We expressed in *N. benthamiana* as GFP fusion proteins either the entire N-terminal region (called Poly-Q-ND) or the Poly-Q domain alone (Fig. 5A). For both constructs, efficient cytosolic foci formation was observed in a HS-dependent manner (Fig. S6). Notably, these constructs displayed a filamentous pattern of the GFP signal that was attenuated under HS and was not observed with full-length GFP-AGO1. Importantly, co-expression of the GFP-Poly-Q together with mRFP-AGO1, confirmed that the HS-induced Poly-Q condensates correspond to those observed with full-length AGO1(Fig. 5C).

To investigate whether the ability to undergo LLPS depends on the properties of the Poly-Q region, we expressed the domain as an N-terminal GFP fusion in *Escherichia coli*, purified it to near homogeneity, and performed *in vitro* phase transition assays. We found that the AGO1 Poly-Q domain, but not maltose binding protein fused to GFP, was sufficient to rapidly drive GFP into droplet-like structures *in vitro* when incubated with the molecular crowding agent PEG 8000 (Fig. 5D-I; Fig. S7A). At near physiological salt concentration, the size of the droplets increased with the PEG concentration from 4 to 10% and their formation was partly inhibited by NaCl concentration above 200mM (Fig. 5D-II). Lower salt concentration was, however, sufficient to initiate the formation of small droplets in absence of PEG (Fig. 5D- I). Droplet size was also strongly influenced by protein concentration, ranging from 1mm to 20mm in diameter when AGO1 Poly-Q domain concentration was raised from 2mM to 20mM (Fig. 5D-III). We next tested whether droplets formation was reversible by diluting ten-fold pre- formed GFP-Poly-Q AGO1 droplets in buffers containing either 1% or 10% PEG. In assays where PEG concentration was reduced to 1%, the droplets mostly disappeared while they remained unchanged when PEG concentration was maintained at 10% (Fig. 5D-IIIV), demonstrating that the process is reversible. To further assess the molecular dynamics and mobility of the AGO1 phase-separated liquid droplets in Arabidopsis, we subjected pAGO1:GFP-AGO1 *ago1-27* line to HS and performed fluorescence recovery after photobleaching (FRAP) assays. In line with the liquid-like nature of the AGO1 droplets, their fluorescence recovery after photobleaching was fast within a few seconds (Fig. 5E). Similar results were obtained when FRAP assays were performed on *in vitro* produced GFP-Poly-Q AGO1 droplets (Fig. S7B).

Since the Poly-Q domain contains several RGG/RG motifs (Xu et al., 2023) recognized as RNA binding motifs (Thandapani et al., 2013), we then investigated whether the presence of RNA had an influence on LLPS. Interestingly, recombinant GFP- Poly-Q domain AGO1 protein could efficiently bind 112nt long single-stranded and double-stranded RNA in electrophoretic mobility assays (EMSA) (Fig. 5F). Moreover, in LLPS assays, when Cy5- labelled RNAs were co-incubated with Poly-Q domain in the presence of 10% PEG, RNA and proteins co-localized in the same droplets (Fig. 5G; Fig. S7C). However, neither droplets size nor droplets abundance were affected suggesting that the interactions between the AGO1 Poly-Q domain and RNA do not facilitate LLPS transition, but only enables co-recruitment in the same structures whose formation is mediated by the protein’s intrinsic properties.

### SGS3 is not required for AGO1 to undergo LLPS and localize in SGs

A previous report showed that RDR6 was not recruited to siRNA bodies in protoplasts of Arabidopsis *sgs3* mutants (Kim et al., 2021), suggesting that the formation of siRNA bodies requires SGS3. To investigate if the formation of condensates containing AGO1 and SGS3 together with SG components during HS also depends on SGS3, we introduced the pAGO1:GFP-AGO1 construct in the *sgs3-14* null mutant (Peragine, 2004). Remarkably, after HS, the formation of AGO1 foci was not impaired in root tip cells, indicating that AGO1 does not rely on SGS3 for LLPS *in vivo* and for its localization to SGs (Fig. 5H). This prompted us to reinvestigate the role of SGS3 for the recruitment of RDR6 during HS in root tip cells of stable transgenic Arabidopsis lines. For this, we analyzed transgenic lines constitutively expressing GFP-RDR6 or RDR6-GFP in both Col-0 and the *sgs3-1* null mutant (Mourrain et al., 2000). After one hr HS at 37°C, RDR6 was able to form foci independent of the presence or absence of SGS3 (Fig. S8), indicating that SGS3 is not essential for the recruitment of RNA silencing components to HS-induced stress granules, at least in Arabidopsis root tip cells.

## DISCUSSION

### During HS AGO1 localizes in SG condensates with siRNA-body, P-body and NMD components

In the present study, we investigated the subcellular distribution of AGO1 in plants subjected to heat stress. Under non-stressed growing conditions, we observed GFP-AGO1 localization mainly diffuse in the cytoplasm as previously reported (Derrien et al., 2012)(Michaeli et al., 2019)(Bologna et al., 2018). Moreover, we did not find an enrichment of the AGO1 signal in any of the endomembrane organelles we examined and that are known to be involved in vesicular trafficking and protein sorting, such as the TGN/EE and MVB. Our observation differs from a study that found that AGO1 is associated with extracellular vesicles (EVs) isolated from the apoplastic fluid of Arabidopsis leaves (He et al., 2021), suggesting AGO1 secretion through the MVB–exosome pathway. However, this contradiction may be explained by the different types of tissues used in the studies: leaf tissue in the former, and root tip cells in our study.

We employed heat stress (HS) to investigate changes in AGO1 subcellular distribution as this stress triggers dramatic effects on gene expression and RNA metabolism (Larkindale and Vierling, 2008)(Merret et al., 2013). We focused our study on three non-membranous RNPs known to play important functions during HS (e.g., P-bodies, SGs and siRNA bodies). In fact, HS blocks translation initiation by provoking 5’-ribosome pausing (Filbeck et al., 2022) and in Arabidopsis, 25% of the paused RNAs undergo 5’-to-3’-mediated decay (Merret et al., 2013). This degradation is mediated by XRN4, an exoribonuclease that resides in P-bodies (Merret et al., 2013). However, paused RNA can also be stored in SGs, which prevent its degradation and allow translation in the recovery period after stress. This is particularly true for mRNA encoding ribosomal proteins (RPs) that are released from SG during recovery for the production of new ribosomes to restore the translation machinery (Merret et al., 2015). Thus, P-bodies and SGs seem to have distinct functions during HS (RNA degradation and RNA storage, respectively).

A third type of condensate, siRNA bodies, contain the essential actors of PTGS, which eliminates dangerous endogenous (transposons) and exogenous (virus) RNA (Bologna and Voinnet, 2014)(Kim et al., 2014). siRNA bodies, composed by the machinery of RDR6/SGS3, form small and discrete cytoplasmic foci which are eventually associated with membranes (Jouannet et al., 2012) (Kumakura et al., 2009). While AGO7, required for the biogenesis of ta-siRNAs from the TAS3 precursor, has been shown to accumulate in siRNA bodies (Jouannet et al., 2012), the relationship of AGO1 with this type of compartment is less understood, despite the fact that AGO1 is a main player in the production of secondary siRNA, including ta-siRNA and pha-siRNA species. Notably, during stress, and particularly HS, it has been shown that siRNA body and SG components co-localize, but not with P-body components (Jouannet et al., 2012; Moreno et al., 2013). Recently, a study reported that a driver of siRNA body formation is SGS3, and the purification of SGS3 condensates identified numerous RNA-binding proteins and components of the PTGS machinery (Tan et al., 2023).

Here we showed that HS triggers a rapid redistribution of AGO1 from its diffuse cytosolic pattern to condensates that also contain siRNA body and SG components. Although microscopy indicated that AGO1 and SGS3 colocalized very well under HS, SGS3 was not found to be significantly enriched via AGO1-immunoprecipitation (IP), and only a few SGS3 peptides were identified by MS. This may be explained by the low abundance of SGS3 protein and its reported potential degradation under HS (Liu et al., 2019). Consistent with this possibility, AGO1 could be co-purified with SGS3 when the latter was overexpressed (Tan et al., 2023). Moreover, both AGO1 (our work) and SGS3 (Tan et al., 2023) interactomes revealed a large number of components of SGs. For instance, we identified PAB8, RNA-binding protein 47A-C (RBP47A-C), RNA helicases (RH6/8/11/52/53), G3BP1, TSN1/2, among others. Nevertheless, we found that although siRNA body and SG components co- localize after HS, these two structures also have distinct features. Indeed, cycloheximide, a drug interfering with translational elongation, inhibits the formation of SGs but not of siRNA bodies. This raises questions about the assembly and dynamics of these entities and whether siRNA bodies remain distinct compartmentalized structures inside SGs.

Intriguing, and less expected, was the identification of some P-body and RNA decay components, such as DCP5 and UPF1, in the AGO1 interactome during HS. We confirmed the colocalization of both proteins with AGO1 during HS in stably transformed Arabidopsis plants. These results indicate that during HS, P-bodies and SGs are not strictly separated entities, but rather that some P body components, and components of nonsense mediated mRNA decay (NMD), can be recruited to SGs. Interestingly, P-bodies and siRNA bodies were often found dynamically juxtaposed, potentially allowing for cross-talk of the two machineries (Martínez de Alba et al., 2015)(Kumakura et al., 2009). At present it is unclear if these proteins are active in the HS-induced condensates, or whether they affect PTGS activity and/or gene expression; however given the central role for UPF1 in posttranscriptional and translational gene regulation (Raxwal et al., 2020), these results deserve further investigation.

### LLPS and recruitment of AGO1 to SG condensates

A large body of previous work revealed that liquid–liquid phase separation (LLPS) of proteins and RNAs are essential for the dynamic assembly of SGs (Molliex et al., 2015)(Protter and Parker, 2016)(Van Treeck et al., 2018). It is also well established that intrinsically disordered regions (IDRs) and prion-like domains (PrLDs) are driving forces for phase separation of many proteins (Shin and Brangwynne, 2017)(Wang et al., 2018). Thus, how condensates are assembled during HS in plants and how AGO1 is recruited to these entities are important questions to answer.

Interestingly, following HS we identified several proteins in the interactome of AGO1 which have the capacity to phase separate into SGs. Among them are ALBA proteins, which undergo LLPS to localize at SGs and were recently shown to confer thermotolerance in Arabidopsis (Tong et al., 2022). Nevertheless, loss of ALBA proteins does not affect the formation of SGs (Tong et al., 2022), indicating that these proteins are not essential for SG assembly. Other proteins identified in our interactome, such as the Tudor Staphylococcal Nuclease (TSN1 and TSN2) and RH8 and RH12, have been shown to contribute to the assembly of SG and P-body condensates (Gutierrez-Beltran et al., 2015)(Chantarachot et al., 2020). Whether these proteins influence the formation of AGO1-containing condensates during HS requires further investigation. Notably, recent work showed that SGS3 exhibits phase separation both *in vivo* and *in vitro* through its PrLDs, highlighting the importance of LLPS in siRNA body formation (Kim et al., 2021)(Tan et al., 2023). These studies also suggested that SGS3 is essential for the recruitment of RDR6 to SGs; in wild-type Arabidopsis protoplasts, RDR6 was able to form siRNA body-like foci, while this was abolished in protoplasts of an *sgs3* mutant (Kim et al., 2021). Moreover, in a yeast heterologous system, it was shown that SGS3-GFP forms condensates and concentrates RDR6-mCherry, while RDR6-mCherry alone is unable to form such entities (Tan et al., 2023). However, in stable Arabidopsis transformants expressing RDR6 fused to GFP, the formation of siRNA body-like foci was still observed under HS, even in a *sgs3* null mutant, indicating that SGS3 is not always required. The discrepancy between our results and the previous reports might be explained by the different experimental settings.

Importantly, our work also revealed that AGO1 does not require SGS3 to efficiently form condensates during HS in Arabidopsis. Interestingly, the still poorly characterized N-terminal region of AGO1 contains a large IDR enriched in polar and charged amino acids, also called Poly-Q, which itself contains a PrLD. Nevertheless, the deletion of the PrLD did not abolish the capacity of AGO1 to form condensates during HS, at least in transient expression assays, indicating that other residues of the Poly-Q contribute to this capacity. The phase-separating behavior of GFP-AGO1 in heat-induced condensates in Arabidopsis cells was also supported by FRAP assays showing a dynamic recovery of the fluorescence after photobleaching. Moreover, the Poly-Q domain alone was sufficient to undergo phase separation both *in vitro* and *in planta*. Strikingly, we found that this domain is also able to bind single-stranded and double-stranded RNA, but the addition of RNA did not facilitate LLPS of the AGO1 Poly-Q domain. It was shown that for some RNA-binding proteins, RNA rather inhibits LLPS to prevent the aberrant formation of protein condensates (Maharana et al., 2018). Whether the Poly-Q domain of AGO1 binds RNA under HS in vivo, and how this would affect AGO1 phase separation, will require further studies. Note that AGO1 may not be the only Arabidopsis AGO protein able to phase separate under HS given the presence of PrLDs in the N-terminal regions of AGO2 and AGO5 (Kim et al., 2021) and the identification of both proteins in the AGO1 interactome during HS. This raises the question of the physiological reasons for the compartmentation of AGO proteins, and for AGO1 in particular, in condensates during stress.

### Effects of a short HS treatment on AGO1 loading and activity

As indicated earlier, P-bodies and SGs have clearly defined function during HS (RNA degradation and RNA storage, respectively). Finding AGO1, but also SGS3 and SG components in the same condensates raises the question of how PTGS is affected by HS and what may be its contribution to the HS response and/or recovery. A previous study reported that the efficiency of transgene PTGS-associated siRNAs and of endogenous tasiRNAs is reduced when growing Arabidopsis plants permanently at 30°C due to the partial dysfunction of the siRNA body component SGS3 (Zhong et al., 2013). Another study reported that the level of Arabidopsis TAS1 ta-siRNAs is reduced and the level of TAS1 ta-siRNA targets increased after one hour of HS at 37°C (Li et al., 2014). Over-expression of TAS1 RNA reduced thermotolerance whereas down-regulation of TAS1 RNA, or over-expression of TAS1 ta- siRNA targets, increased thermotolerance. Conversely, a recent study (Tan et al., 2023) indicated that the liquid-like properties of SGS3 are essential for its function in siRNA production (Tan et al., 2023). The authors reported that 15 min of HS at 42°C or treatment with cycloheximide (CHX), which both inhibit translation, increases the level of ta-siRNAs and causes the production of siRNAs from endogenous genes; similar outcomes are observed when P-body components are impaired suggesting that siRNA bodies are active during a short HS treatment (Tan et al., 2023). On the contrary, our results, from WT Col-0 seedlings subjected to 1 hr HS at 37°C, revealed that although the transcriptome strongly responds to HS, changes at the sRNA level are quite minor. In particular, we did not observe a strong accumulation of 21/22 nt siRNAs from endogenous genes nor a significant increase in any of the tasiRNA population. Further, the relative abundance of most miRNA, or their loading in AGO1, were not significantly affected by the HS treatment. However, this does not exclude that some of the miRNAs could play a function during HS, as a number of unique miRNA target signatures were detected during HS and recovery. The discrepancy between our data and the recent report (Tan et al., 2023), could be explained by the fact that the latter study used SGS3- overexpressing approaches, while we used WT Col-0 seedlings. It is also possible that the level of temperature and/or the length of the HS modulates siRNA production and PTGS activity. Indeed, when focusing only on tasiRNA accumulation, a permanent growth at 30°C resulted in reduced accumulation (Zhong et al, 2013), whereas 1 hr at 37°C caused limited (Li et al, 2014) or no significant reduction (our work), contrasting a shorter but more intense HS (15 min at 42°C), which caused an increased accumulation (Tan et al., 2023). Overall, understanding the impact of increased temperatures on AGO1 and more globally on RNA silencing will be crucial to develop novel strategies to cope with climate change and global warming.

## SUPPLEMENTAL INFORMATION

Supplemental information can be found online

## ACKNOWLEDGMENTS

We thank Olivier Voinnet for the pAGO1:GFP-AGO1 *ago1-27* and pAGO1:mCherry-AGO1 lines. P.G. acknowledges support from the European Research Council under the European Union’s Seventh Framework Programme (FP7/2007-2013) / ERC advanced grant to PG agreement n° [338904] and Agence Nationale de la Recherche (ANR) grant [HEAT-STRESS- BODIES, ANR-23-CE11-0014-02] and IdEx Unistra (ANR-10-IDEX-0002), SFRI-STRAT’US project (ANR-20-SFRI-0012) and EUR IMCBio (IMCBio ANR-17-EURE-0023). The mass spectrometry instrumentation was funded by the University of Strasbourg, IdEx “Equipement mi-lourd” 2015.

## AUTHOR CONTRIBUTIONS

AB and PG designed research; AB, PB, MS, EL, NB, PH, TE, HV performed research; AB, PB, MS, EL, NB, PH, TE, DG, HV, BM and PG analyzed the data; PG wrote the paper, with help from AB, PB and HV.

## Materials availability

Transgenic plant seeds generated in this study are available from the Lead Contact on request.

## Data and Code Availability

Availability of proteomics and RNA-seq data through online repertories: The mass spectrometry proteomics data have been deposited to the ProteomeXchange Consortium via the PRIDE (Perez-Riverol et al., 2022) partner repository with the dataset identifier PXD044228 and 10.6019/PXD044228. The deep sequencing data have been deposited in NCBI’s Gene Expression Omnibus (Edgar et al., 2002) and are accessible through GEO Series accession number GSE239837.

## EXPERIMENTAL PROCEDURES

### Experimental model

*Arabidopsis thaliana* ecotype Colombia as well as *Nicotiana benthamiana* (for transient expression assays), were used in this study. The Arabidopsis mutants, *ago1-27* (Morel et al., 2002) and the *sgs3-14* null mutant (Peragine, 2004) were used. All transgenic Arabidopsis lines are detailed in Table S1. The list of primers used for genotyping is presented in Table S3.

### Plasmid constructions

The list of primers used for cloning is presented in Table S3.

pENTRY(221)-Poly-Q: To obtain this construct, the N-terminal sequence of AGO1 (CDS) corresponding to the Poly-Q domain of AGO1 (amino acid 1 to 207) was amplified from the vector pENTRY(ZEO)-AGO1 (CDS) with oligo primers containing AttB1 and AttB2 recombination sites (primers called AGO1 Poly-Q-fwd and AGO1 Poly-Q-rev, see Table S3) and the sequence was mobilized into the pDONR221 vector by BP Gateway recombination (Invitrogen).

pENTRY-R2-Poly-Q-ND-L3: To obtain this construct, the N-terminal sequence of AGO1 corresponding to the Poly-Q-ND domain of AGO1 (amino acid 1 to 300) was amplified from the vector pENTRY-R2-AGO1genomic-L3 with oligo primers containing AttB3 and AttB2R recombination sites (primers called attB2R_AGO1g_F and attB3NterAGO1-300_rev, see Table S3) and the sequence was mobilized into the pDONOR-P2RP3 vector by BP Gateway recombination (Invitrogen).

The pENTRY(ZEO)-AGO1(DPrLD) vector has been obtained by mutagenesis using pENTRY(ZEO)-AGO1(CDS) as a template by the company GenScript. In this construct a sequence of 119 amino acids (amino acid 13 to 132) encompassing the PrLD domain has been deleted from WT AGO1 (CDS).

The p35S:GFP-Poly-Q construct was obtained by Gateway LR recombination (Invitrogen) using pENTRY(221)-Poly-Q AGO1 and the binary vector pB7WGF2 ((Karimi et al., 2005); https://gatewayvectors.vib.be/collection). This construct expresses a GFP-Poly-Q-AGO1 fusion protein under the regulation of the 35S promoter.

The p35S:GFP-Poly-Q-ND construct was obtained by assembling the 35S promoter (pENTRY-R4-35S-L1), the GFP (pENTRY(221)-GFP) and the Poly-Q-ND sequence of AGO1(pENTRY-R2-Poly-Q-ND-L3) into the binary vector pB7m34GW (https://gatewayvectors.vib.be/collection) by the three-way LR Gateway reaction (Invitrogen).

This construct expresses the GFP-Poly-Q-ND of AGO1 fusion under the control of the 35S promoter.

The p35S:GFP-AGO1(DPrLD) construct was obtained by Gateway LR recombination (Invitrogen) using pENTRY(ZEO)-AGO1(DPrLD) and the binary vector pB7WGF2 ((Karimi et al., 2005); https://gatewayvectors.vib.be/collection). This construct expresses the GFP- AGO1(DPrLD) fusion protein placed under the regulation of the 35S promoter.

The p35S:mRFP-AGO1 construct was obtained by Gateway LR recombination (Invitrogen) using pENTRY(ZEO)-AGO1(CDS) (Hacquard et al., 2022) and the binary vector pB7WGR2 (Karimi et al., 2005); https://gatewayvectors.vib.be/collection. This construct expresses the mRFP-AGO1(CDS) fusion protein under the regulation of the 35S promoter.

The pRDR6:SGS3-GFP construct was obtained by a two-step process. First pRDR6-pGWB4 was obtained by cloning RDR6 promoter in pGWB4 (Nakagawa et al., 2007) digested by HindIII after amplification by the primers pairs proSGS2F-hIII / proSGS2R-HIII (see Table S3). In parallel the cDNA of *SGS3* was cloned in the GATEWAY™ compatible vector pDONR207 (Invitrogen) using the following primers: attB2SGS3f/ attB2SGS3R (see Table S3). Finally, SGS3 was transferred to the binary vectors pRDR6-pGWB4 by Gateway LR recombination (Invitrogen) to make the pRDR6:SGS3-GFP.

### Plant growth conditions and treatments

For *in vitro* culture conditions, Arabidopsis seeds were surface-sterilized using ethanol and plated on MS agar (MES-buffered MS salts medium [Duchefa, Murashige & Skoog medium inc. vitamins/MES- MO255)], 1% sucrose, and 0.8% agar, pH 5.7). The seeds were then stratified for 2 days at 4°C in the dark and then transferred in 16h light/8h dark (20,5/17°C,70% humidity) growth chamber, under fluorescent light (Osram Biolux 58W/965). The plants that grew on soil were under a 16h light/8 h dark diurnal regime.

For root cell microscopy, seeds were grown on MS-agar plates that were positioned vertically in the growth chamber.

For HS treatment, 7-day old seedlings were mounted on microscopy slides, that were transferred into an incubator set at 37°C for 30 minutes.

For cycloheximide treatments, Arabidopsis seedlings were grown for 5 days on MS-agar plates then transferred into liquid 1/2MS medium, 0,5g/L sucrose supplemented with either 100µM of cycloheximide (CHX) or methanol (0,02%) (Mock) for 30 minutes. These seedlings were then mounted on microscopy slides and put in an incubator heated at 37°C for 30 minutes.

For western blot, qRT-PCR, Northern blot and IP-MS experiments, Col-0 and pAGO1:GFP- AGO1 *ago1-27* lines were grown for 7 days on MS-agar plates then exposed to 37°C for 1 hour. In order to get the most homogeneous heat treatment, the MS-agar plates were wrapped into a plastic bag and submerged for 1 hour into a water bath set at 37°C. The plates were transferred into a second water bath set at 22°C for 2hours when recovery experiments were performed.

### Transient expression in N. benthamiana leaves

Agrobacterium cells (GV3101 Pmp90 or C58C1) harboring the constructs of interest were grown overnight at 28°C in 10mL LB medium supplemented with antibiotics, resuspended in 10mM MgCl_2_ supplemented with 200mM acetosyringone at an OD of 0,3 per construct (unless otherwise specified), and incubated for 1 hour at room temperature before being pressure infiltrated into leaves of 4 week-old plants. All agro-infiltration assays were conducted in the presence of P19. Plants were maintained in growth chambers under 16 hours light and 8 hours dark photoperiod with a constant temperature of 22°C. Sampling of leaf disc was performed using a mechanical sampler and observations of 3 leaf discs, each coming from a different leaf and plant, were performed 48 hours after agro-infiltration. Agroinfiltrated leaves were heat stressed by incubation in a previously heated at 37°C 10mM MgCl_2_ buffer for 1 hour at 37°C in darkness.

### Confocal microscopy analysis, quantifications and statistical analysis

Tobacco leaf (abaxial epidermal cells) and Arabidopsis root cells were imaged by CLSM using a Leica SP8 microscope. Fluorescence Recovery After Photobleaching (FRAP) assays were performed using a Zeiss LSM780 microscope. Arabidopsis seedling roots were positioned between a microscopy slide and a cover slip containing 0.5MS 0.5g/L sucrose liquid media, while hypocotyls and cotyledons were left exposed. Usual excitation/detection-range parameters for GFP and mCherry, mRFP or tagRFP were 488 nm/500–550 nm and 561 nm/600–650 nm, respectively and emissions were collected using the system’s hybrid (Hyd) and photomultiplicator detectors. When GFP and mRFP/mCherry/tagRFP were simultaneously imaged in transient expression assays, excitation/detection-range parameters were 488 nm/500–550 nm and 561 nm/600–650 nm, respectively. Sequential scanning was employed at all times. Images were processed and analyzed using ImageJ.

Quantification of GFP or RFP labeled bodies was done semi-automatically using the ImageJ macro termed “2CH_bar” described previously (Michaeli et al., 2019). In each sample, density of foci per cell was calculated by first measuring the surface size covered by GFP signal (in µm2) and dividing the number of foci by the GFP surface size (number of foci/µm2). The 2CH_bar macro also allowed to determine the degree of correlation between green (GFP/YFP) and red (mCherry/tagRFP/mRFP) tagged bodies. For all samples and in each channel, data (XY coordinates and relative green/red fluorescence intensity) were gathered on each tagged body recognized by the macro which indicates also the total number of foci spotted. Then, global Pearson Coefficient Calculation was performed for each channel using relative green/red fluorescence intensity obtained for each spotted dot. Data are presented as mean ± SEM.

### Protein analysis and western blotting

Proteins were extracted in pre-heated (95°C) 2X Laemmli sample buffer (62 mM Tris HCl pH 6.8, 3% SDS, 40% glycerol, 0,1% bromophenol blue, DTT 100mM, quantified using amido- black staining (Popov et al., 1975) and 20µg of total proteins were separated by SDS-PAGE, either on 7–12% Tris-glycine gels or gradient NuPAGE 4–12% Bis-Tris Protein Gels (Thermo Fischer) or gradient Criterion TGX gel (4–15%) (BioRad). A list of antibodies and their working dilution used in this work are reported in (Table S3). For all western blots, immunoluminescence was detected using the ECL Clarity (BioRad) and imaged using Fusion FX (Vilbert)

### Protein immuno-precipitation assays

For each GFP-IP, 1g of seedlings was ground in liquid nitrogen for 10 minutes in 3ml of ice-cold lysis buffer (50mM Tris-HCl pH7,5, 50mM NaCl, 0,25% IGEPAL CA-630, 2mM MgCl2, 5mM DTT, protease inhibitors (Complete™–EDTA free, Roche). The supernatant was cleared by 2 centrifugation steps of 15min then 5min at 10,000g at 4°C. The cleared supernatants were divided in 3 affinity purifications, incubated with magnetic microbeads coupled to GFP antibodies (µMACS™ GFP Isolation Kit, Miltenyi Biotec) and complexes were eluted in 100µl of pre-warmed elution buffer. IP experiments were performed in four independent biological replicates. Each biological replicate was divided into three affinity purification replicates. In parallel control IPs were carried out with GFP antibodies in Col-0.

For immunoprecipitation of endogenous AGO1, 1g of frozen tissues was ground to a fine powder with a mortar and pestle, resuspended in 3 volumes of crude extract buffer (50mM Tris, pH 7,5, 150mM NaCl, 10% glycerol, 5mM MgCl2, 0,2% IGEPAL, 5mM DTT, and 1x Complete™–EDTA free (Roche)) and incubated for 20 min at 8 rpm in the cold room. Insoluble material was removed by centrifugation (two centrifugation steps at 10,000g at 4°C, first for 15 min then for 5 min). Identical amounts of crude extracts were incubated with prebound @AGO1 (5µg of @ AGO1 from Agrisera) PureProteome Protein A magnetic beads (30µL; Millipore) for 2 hr at 7 rpm at 4°C. Immune complexes were washed four times in the crude extract buffer, and purified sRNA was eluted from the beads in Tri-Reagent (Sigma-Aldrich) following the manufacturer’s instructions. Extracted RNA was precipitated in 2 volumes of isopropanol + 40 μg glycogen overnight at −20°C. RNA was resuspended in sterile water.

### Mass spectrometry analysis, data processing and availability

Eluted proteins were digested with sequencing-grade trypsin (Promega, Fitchburg, MA, USA). Each sample was further analyzed by nanoLC-MS/MS on a QExactive+ mass spectrometer coupled to an EASY-nanoLC-1000 (Thermo-Fisher Scientific, USA) as described previously (Chico et al., 2020). Data were searched against the TAIRv10 fasta protein sequences from *Arabidopsis thaliana* with a decoy strategy (27.282 forward protein sequences). Peptides and proteins were identified with Mascot algorithm (version 2.8, Matrix Science, London, UK) and data were further imported into Proline v2.0 software (http://proline.profiproteomics.fr/). Proteins were validated on Mascot pretty rank equal to 1, and 1% FDR on both peptide spectrum matches (PSM score) and protein sets (Protein Set score). The total number of MS/MS fragmentation spectra was used to quantify each protein from at least six independent biological and affinity replicates. After a DEseq2 normalization of the data matrix, the spectral count values were submitted to a negative-binomial test using an edgeR GLM regression through R (R v3.2.5). For each identified protein, an adjusted pvalue (adjp) corrected by Benjamini–Hochberg was calculated, as well as a protein fold-change (FC). The results are presented in a Volcano plot using protein log2 fold changes and their corresponding adjusted p-value (-log10adjp) to highlight enriched and depleted proteins.

### Total RNA extraction

Total RNAs were extracted from 7-d-old seedlings grown on MS agar plates using TRI Reagent (Sigma) following the manufacturer’s instructions. RNAs were precipitated O/N at -20°C in 2 volumes of isopropanol + 40 μg glycogen. Quality and concentration of purified RNAs were assessed using Nanodrop and Qubit fluorometer.

### RT-qPCR

For RT-qPCR, 2 μg of total RNA treated with DNase RQ1 (PROMEGA) was reverse transcribed with Superscript IV (Invitrogen) using a mix of random hexamers and oligo d(T).

Each quantitative PCR reaction was performed in three technical replicates with Takyon™ SYBR® 2X qPCR Mastermix Blue (Eurogentec) in 384-wells plates with a total volume of 10 uL using Light Cycler 480 apparatus (Roche). mRNA abundance was compared to two reference genes EXP (AT4G26410) and TIP41 (AT4G34270) (Table S3). mRNA relative abundance was calculated using the Λ1Λ1Ct method.

### Low molecular weight northern blot

Northern blot analyses of low molecular weight RNAs were performed with 15 μg of total RNA resuspended in a final concentration of 50% (v/v) formamide-5mM EDTA-0.05% (w/v) bromophenol blue-0.05% (w/v) xylene cyanol, heated at 95°C for 2 min, and separated by electrophoresis on 15% polyacrylamide gels (19:1 acrylamide:bisacrylamide), 8 M Urea, 0.5× TBE gel. RNA was then transferred on Hybond-NX (Amersham) membrane and crosslinked with EDC for 1h30 at 60 °C. DNA oligonucleotides complementary to sRNA and U6 were end-labeled with [γ-32P]ATP using T4 PNK (ThermoFisher). Membranes were incubated at 42 °C with PerfectHyb Plus hybridization buffer (Sigma-Aldrich) for 30 min, hybridization was performed overnight in PerfectHyb Plus containing the radiolabelled probe at 42 °C. The sequences of the Northen blot probes are described in Table S3). Membranes were washed three times in 1× SSC-1% SDS before exposure to a Fujifilm imaging plate. Signals were visualized with an Amersham Typhoon IP Biomolecular Imager (GE Healthcare Life Sciences).

### ssRNA and dsRNA substrate preparation

A 112 nt long product from the GFP sequence was PCR amplified from plasmid pB7WGF2 with two oligos pairs (A349/A347 and A346/A348) using Phusion polymerase (ThermoFisher). The PCR products were purified on gel, and used as template in two separate T7 polymerase *in vitro* transcription reactions to generate sense and antisense transcripts. For Cy5 labelled ss and dsRNA, the *in vitro* transcription reaction of the sense transcript was spiked with 1mM Cy5-allyl-UTP (Jena Bioscience). RNA transcripts were purified from the reaction with the RNA clean-up Kit (Mascherey-Nagel), quantified with a Qubit fluorometer (ThermoFisher) and verified on agarose gel. Double stranded RNA was obtained by equimolar mixing and annealing of both reverse-complementary transcripts.

### Recombinant polyQ-AGO1 and MBP domain expression and purification

The Arabidopsis AGO1 DNA sequence encoding the prion-like motifs and intrinsically disordered regions of the N-terminal domain (amino acid 2 to 207) was amplified by PCR from a previously validated cDNA with two pairs of oligos (AGO int R, AGO1int F, AGO1-pQ B4 F and AGO1-pQ B4 R, see Table S3) in order to delete an internal *BsaI* site and generate a sequence compatible with GoldenGate cloning. The amplified products were digested, re- ligated within the *BsmBI* sites of the pUPD2 (Sarrion-Perdigones et al., 2014) vector before being further assembled into the *BsaI* sites of a customized GoldenGate-compatible pET expression plasmid to generate the final GFP-AGO1pQ domain-6xHis construct under the regulation of a T7 promoter. The GFP-MBP control plasmid was assembled similarly with the E.coli Maltose-binding-protein sequence replacing polyQ-AGO1 sequence to generate GFP-MBP-6xHis.

GFP-Poly-Q-6xHis and GFP-MBP-6xHis expressions were realized in BL21 *E. coli* strain in auto-inducing medium (Studier 2005) at 20°C for 18h. Cells were collected by centrifugation, resuspended in 25mM HEPES pH7.4, 500mM NaCl, 5mM beta-mercaptoethanol, and lyzed by three passages on a microfluidizer LM-20 (Microfluidics) at 1200 bars at 4°C. The lysates were clarified by centrifugation at 18000g for 30min at 4°C and the supernatants loaded on a NiNTA affinity column (HisTrap FF crude, Cytiva) equilibrated in 25mM HEPES pH7.4, 500mM NaCl, 25mM Imidazole and eluted in the same buffer with 250mM imidazole. The proteins were further purified through a size exclusion chromatography Superdex 200 16/60 column (Cytiva) equilibrated in 250mM HEPES pH7.4, 500mM NaCl. Fractions corresponding to monomeric proteins (apparent MW of 130 kDa for GFP-polyQ protein and 100kDa for GFP-MBP protein) were collected and concentrated by ultrafiltration, flash frozen in liquid nitrogen and stored at -80°C until use. Quality of proteins were assessed on 12% Tris-glycine SDS-PAGE gel, stained with Coomassie blue. Quantification was done by UV absorbance at 280nm on a Nanodrop 2000 (Thermo Scientific).

### In vitro phase separation assay

For *in vitro* liquid droplet formation, purified protein was brought to the desired concentration through dilution into the reaction buffer containing 25mM HEPES pH 7.5, 200mM NaCl and 10% PEG 8000 (unless specified otherwise). 5ul droplets were placed on a glass slide and sealed in a small chamber, to prevent evaporation. After a 15min incubation period, imaging was performed under an LSM-700 Zeiss confocal microscope using 20X or 40X objectives. For co-localization experiment, 10x concentrated Cy5-labelled single stranded and double-stranded RNAs were added to the reaction together with the protein and the PEG containing dilution buffer, mixed thoroughly and incubated for 15min before observation. GFP and Cy5 were excited at 488 and 633nm, respectively, and detected at 490-612nm and 638-759nm, respectively.

### FRAP assays

FRAP analyses were performed on an LSM-780 Zeiss using a 40X objective on samples prepared as specified above. For *in vivo* FRAP assays, photobleaching was achieved with the laser set at 488nm and 405nm, at 40% and 17,6% intensity, respectively, and 140 iterations). For *in vitro* FRAP assays photobleaching was achieved with the laser set at 488nm, 40% intensity and 120 iterations).

### EMSA

Recombinant GFP-polyQ AGOI protein and *in vitro* transcribed RNAs were produced as described above, except that RNA was not fluorescently labeled. 9pmol of RNA was mixed with the specified amount of protein (5,6-20pmol by 2-fold increase) in 10ul volume reaction containing HEPES 25mM pH7.5, NaCl 50mM. After incubation at 25°C for 15min, the reaction was resolved on a 1% agarose gel, 0,5X Tris Borate EDTA buffer containing 1ug/ml ethidium bromide, at 100V, during 45min at RT. Gel documentation was done in a Fusion FX system, with the excitation at 365nm and the emission signal recorded at 595nm.

### Libraries preparation and high-throughput sequencing

Total RNA samples were extracted from 1-week-old Col-0 seedlings (Col-0 seedling +/- 1hour treatment at 37°C (HS) and after an additional 2H period back at 22°C (Recovery)) grown on MS-agar plates using Tri-Reagent according to the manufacturer’s instruction. For AGO1- loaded sRNA samples, AGO1-IPs were performed as described above from 1g of 1-week-old Arabidopsis seedlings grown on MS-agar plates and following similar HS and recovery treatments. AGO1-loaded sRNAs were then extracted by adding Tri-Reagent directly on the magnetic beads and extraction of RNA was performed according to the manufacturer’s instructions. RNAs were precipitated O/N at -20°C in 2 volumes of isopropanol + 40 μg glycogen as described above.

Small RNA seq libraries were generated using RealSeq v2 kit (Barberán-Soler et al., 2018), following manufacturer’s instructions, with 100ng as initial input and 15 cycles PCR amplification. nanoPARE and RNAseq libraries were generated following the protocol published by (Schon et al., 2018). All libraries were sequenced using Illumina NextSeq technology at the Delaware Biotechnology Institute (DE, USA). For small RNA libraries, we trimmed adapters and low-quality reads using Cutadapt v2.9 software (Martin, 2011) and retaining only reads between 18- and 34-nt long. Reads were then mapped to the Arabidopsis genome version 10 (available at www.arabidopsis.org/download/) and its corresponding TAIR10 blastsets for all the features, using Bowtie2 (Langmead and Salzberg, 2012). NanoPARE libraries were analyzed using the pipeline provided by the authors (http://www.github.com/Gregor-Mendel-Institute/nanoPARE). RNAseq libraries were analyzed unsing the HISAT2 and StringTie pipeline (Pertea et al., 2016). Differential accumulation and expression were done using DESeq2 (Love et al., 2014) and all plots were generated using ggplot2 (Wickham, 2010) packages in R environment.

Additional information for the sequencing experiments is presented in Tables S4-S7. Table S4: Summary table for all sequencing experiments; Table S5: RNAseq summary table; Table S6: Small RNA count summary; Table S7: Nano-PARE summary.

## QUANTIFICATION AND STATISTICAL ANALYSIS

Barplot in Fig. 1C, Volcano plot in Fig. 3A and the line chart in Fig. 4A have been generated with R version 3.6.3, running under macOS Sierra 10.12.6. Packages used for the volcano plot were: ggplot2 (v3.4.1), dplyr(v1.1.1) and ggrepel (v0.9.2). Additional packages used for barplot and boxplot were gridExtra(v2.3) and cowplot(v1.1.1).

---

**Fig. S1.**
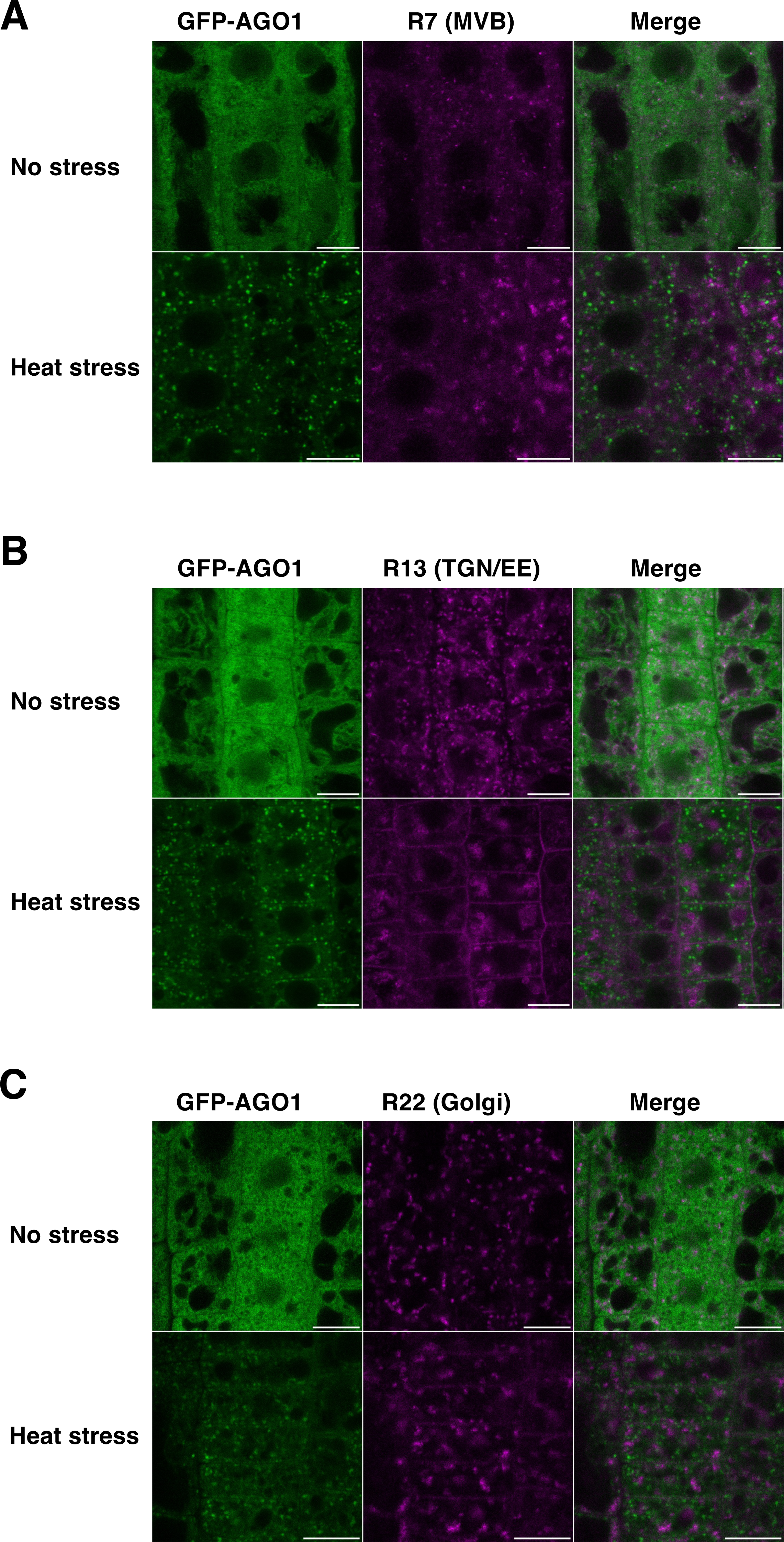
Dual subcellular localization of AGO1 with different cytoplasmic compartment markers. **(A)** AGO1 does not colocalize with the MVB marker R7 in root tip cells. CLSM on 5-day old Arabidopsis pAGO1:GFP-AGO1 *ago1-27* x pUB10:mCherry-Rha1/RabF2a (R7) root tip cells before and after 37°C HS of 30 minutes. Experiments were performed in triplicate with three roots per replicate. Bar = 10 µm. Foci number before HS for R7 = 1451 ± 308; PCC= 0.34 ± 0.13 Foci number after HS for R7 = 894 ± 310; PCC= 0.29 ± 0.15 Foci number after HS for GFP-AGO1 = 1525 ± 475; PCC= 0.19 ± 0.10 **(B)** AGO1 does not colocalize with the TGN/EE marker R13 in root tip cells. CLSM on 5 day old Arabidopsis pAGO1:GFP-AGO1g *ago1-27* x pUB10:mCherry-VTI12 (R13) root tip cells. Before and after 37°C HS of 30 minutes. Experiments were performed in triplicate with three roots per replicate. Bar = 10 µm. Foci number before HS for R13 = 1311 ± 478; PCC= 0.12 ± 0.17 Foci number after HS for R13 = 553 ± 355; PCC= 0.07 ± 0.08 Foci number after HS for GFP-AGO1 = 1370 ± 438; PCC= 0.04 ± 0.09 **(C)** AGO1 does not colocalize with the Golgi marker R22 in root tip cells. CLSM on 5 day old Arabidopsis pAGO1:GFP-AGO1 *ago1-27* x pUB10:mCherry-SYP22 (R22) root tip cells before and after 37°C HS of 30 minutes. Experiments were performed in triplicate with three roots per replicate. Bar = 10 µm. Foci number before HS for R22 = 1355 ± 508; PCC= 0.47 ± 0.11 Foci number after HS for R22 = 662 ± 355; PCC= 0.28 ± 0.17 Foci number after HS for GFP-AGO1 = 1157 ± 751; PCC= 0.27 ± 0.15

**Fig. S2.**
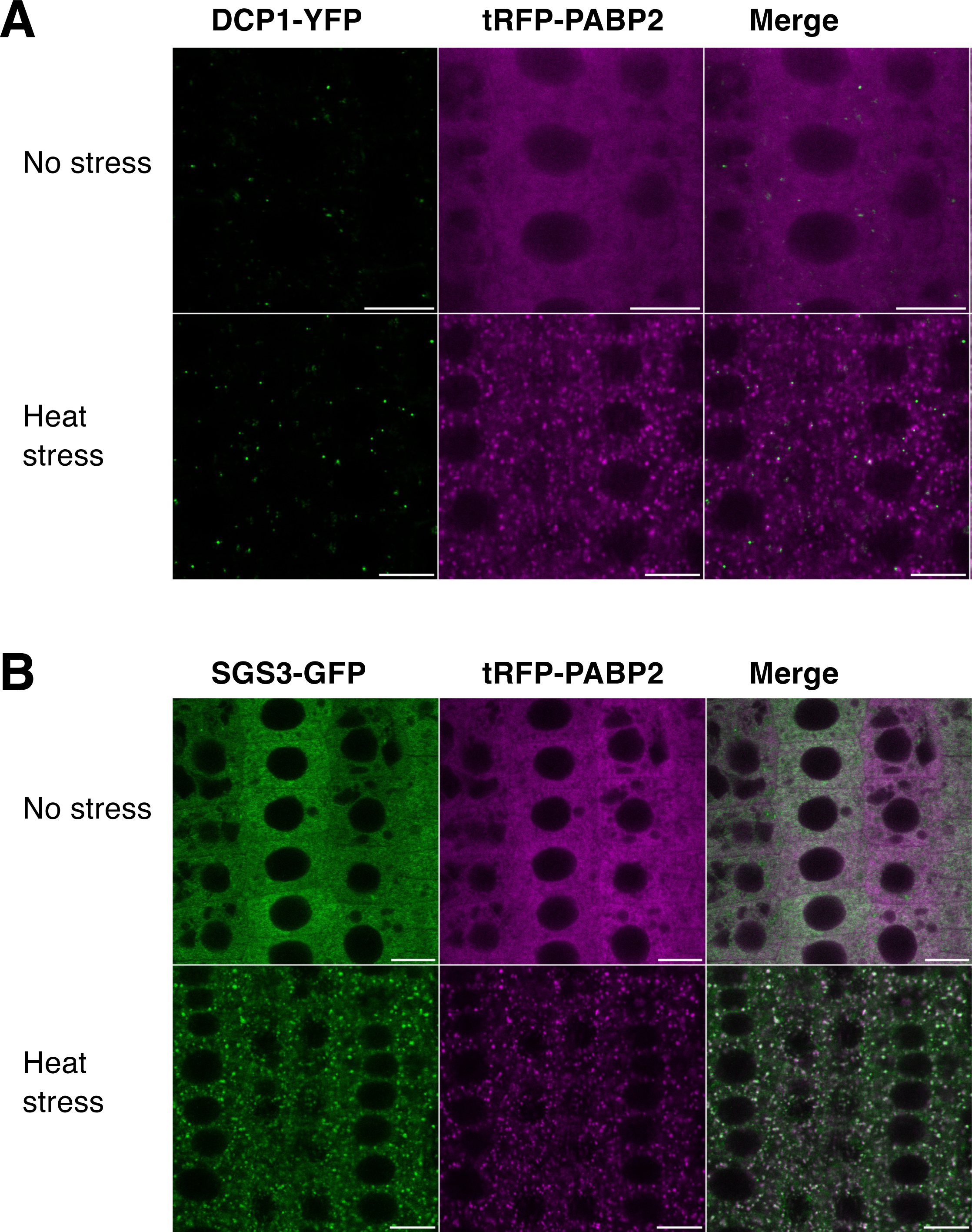
The stress granule marker PAB2 colocalizes in foci with SGS3, but not DCP1 under HS. **(A)** CLSM imaging of 5 day old Arabidopsis pDCP1:DCP1-YFP *dcp1-3* x pPABP2:tRFP-PABP2 root tip cells before and after 37°C HS of 30 minutes. Experiments were performed in triplicate with three roots per replicate. Bar = 10 µm. Objective 40X, oil immersion. Foci number before HS for DCP1-YFP = 1300 ± 458; PCC= 0.15 ± 0.10. Foci number after HS for DCP1-YFP = 478 ± 150; PCC= 0.06 ± 0.15. Foci number after HS for tRFP-PABP2 = 1666 ± 395; PCC= 0.34 ± 0.13. **(B)** CLSM imaging of 5 day old Arabidopsis pRDR6:SGS3-GFP x pPABP2:tRFP-PABP2 root tip cells before and after 37°C HS of 30 minutes. Experiments were performed in triplicate with three roots per replicate. Bar = 10 µm. Foci number after HS for SGS3-GFP = 4156 ± 1310; PCC= 0.83 ± 0.05. Foci number after HS for tagRFP-PABP2 = 6458 ± 1787; PCC= 0.83 ± 0.04.

**Fig. S3.**
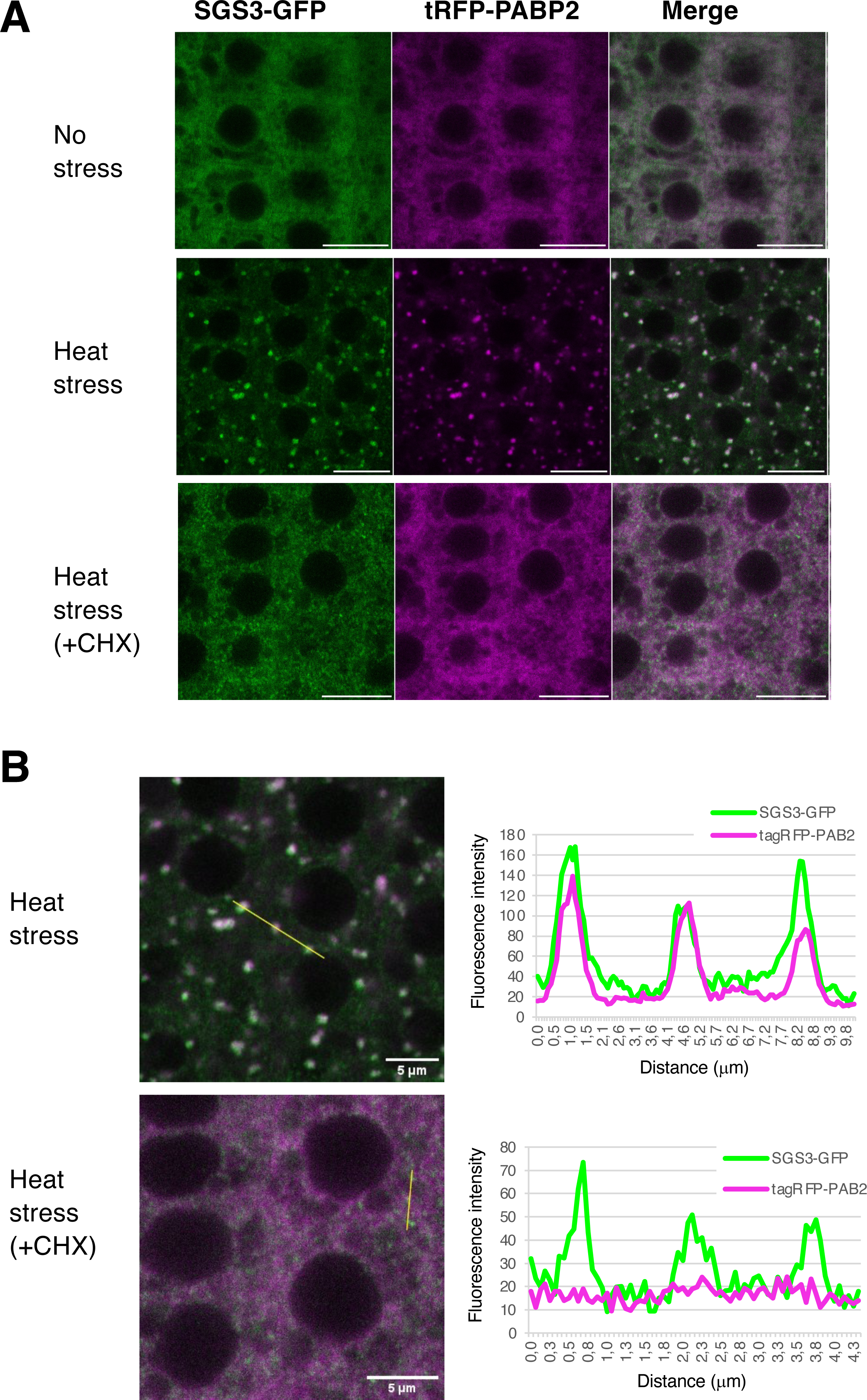
Cycloheximide inhibits stress granule formation, but not small SGS3 foci. (A) CLSM imaging of 5 day old Arabidopsis pRDR6:SGS3-GFP x pPABP2:tRFP-PABP2 root tip cells before HS (upper panels), after 37°C HS of 30 minutes (middle panels) and after 37°C HS of 30 minutes in the presence of 100µM CHX). Experiment performed in duplicate with three roots per replicate. Bar = 10 µm (B) Signal intensity distribution of the total amount of pixels at the x axis shown in the CHX- untreated and CHX-treated cells after 30 minutes of HS at 37°C shown in A. Scale bar = 5 µm. Colocalization of SGS3-GFP and tRFP-PABP2 in CHX-untreated cells: PCC = 0.94 Colocalization of SGS3-GFP and tRFP-PABP2 in CHX treated cells: PCC = 0.18

**Fig. S4.**
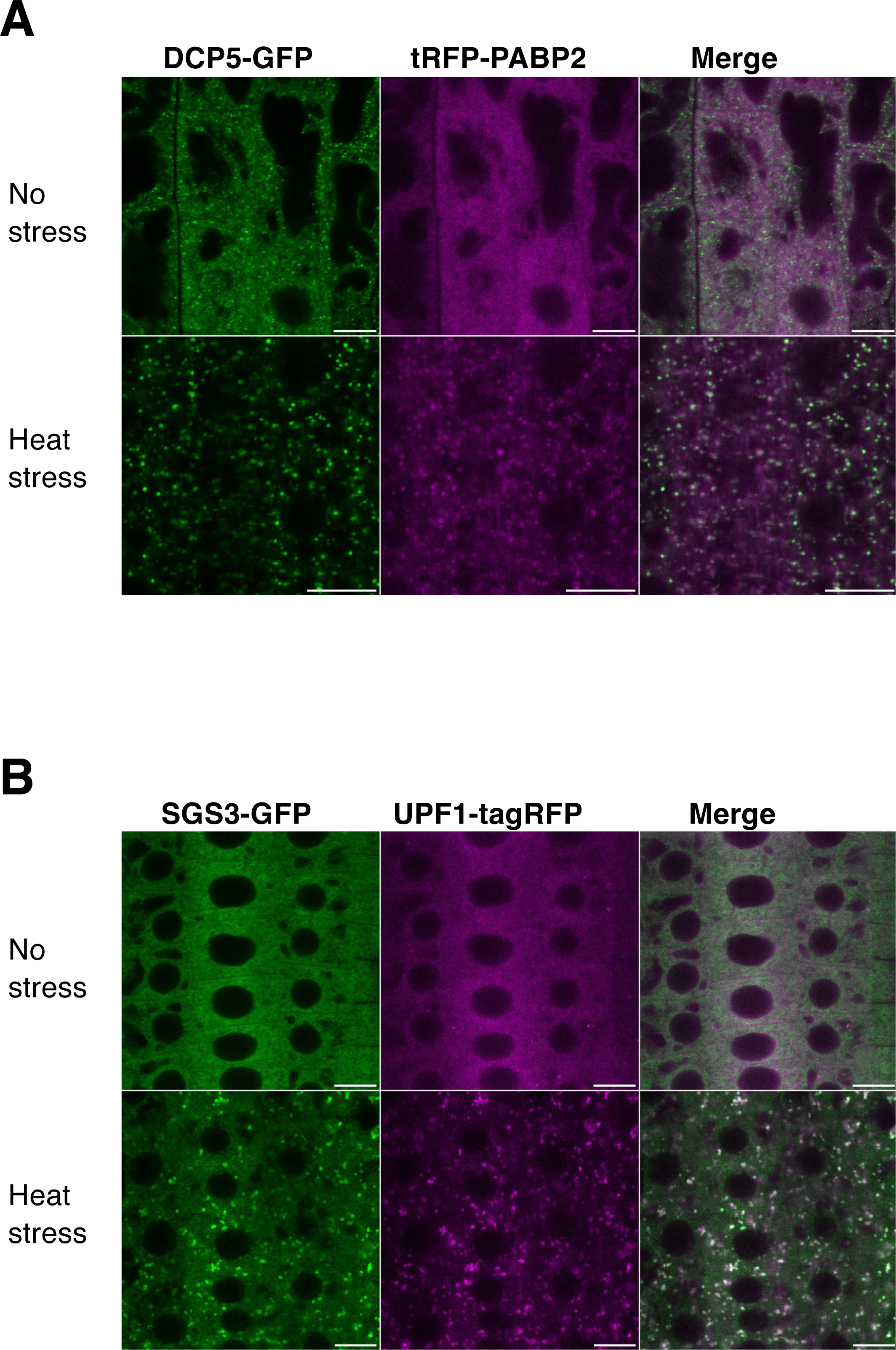
DCP5/PAB2 and SGS3/UPF1 colocalization during HS. **(A)** DCP5 colocalizes with PAB2 in foci under heat-stressed conditions. Confocal laser scanning microscopy on 5 day old Arabidopsis pUB10:DCP5g-GFP *dcp5-1* x pPABP2:tRFP-PABP2 root tip cells before and after 37°C HS of 30 minutes. Experiments were performed in triplicate with three roots per replicate. Bar = 10 µm. Foci number after HS for DCP5-GFP = 479 ± 128; PCC= 0.67 ± 0.11. Foci number after HS for tRFP-PABP2 = 400 ± 37; PCC= 0.61 ± 0.12. **(B)** SGS3 colocalizes with UPF1 in foci under heat-stressed conditions. Confocal laser scanning microscopy on 5 day old Arabidopsis pRDR6:SGS3-GFP x pUPF1:UPF1g-tagRFP *upf1-5* root tip cells before and after 37°C HS of 30 minutes. Experiments were performed in triplicate with three roots per replicate. Bar = 10 µm. Foci number after HS for SGS3-GFP = 1498 ± 641; PCC= 0.77 ± 0.04 Foci number after HS for UPF1-tagRFP= 1602 ± 613; PCC= 0.77 ± 0.05

**Fig. S5.**
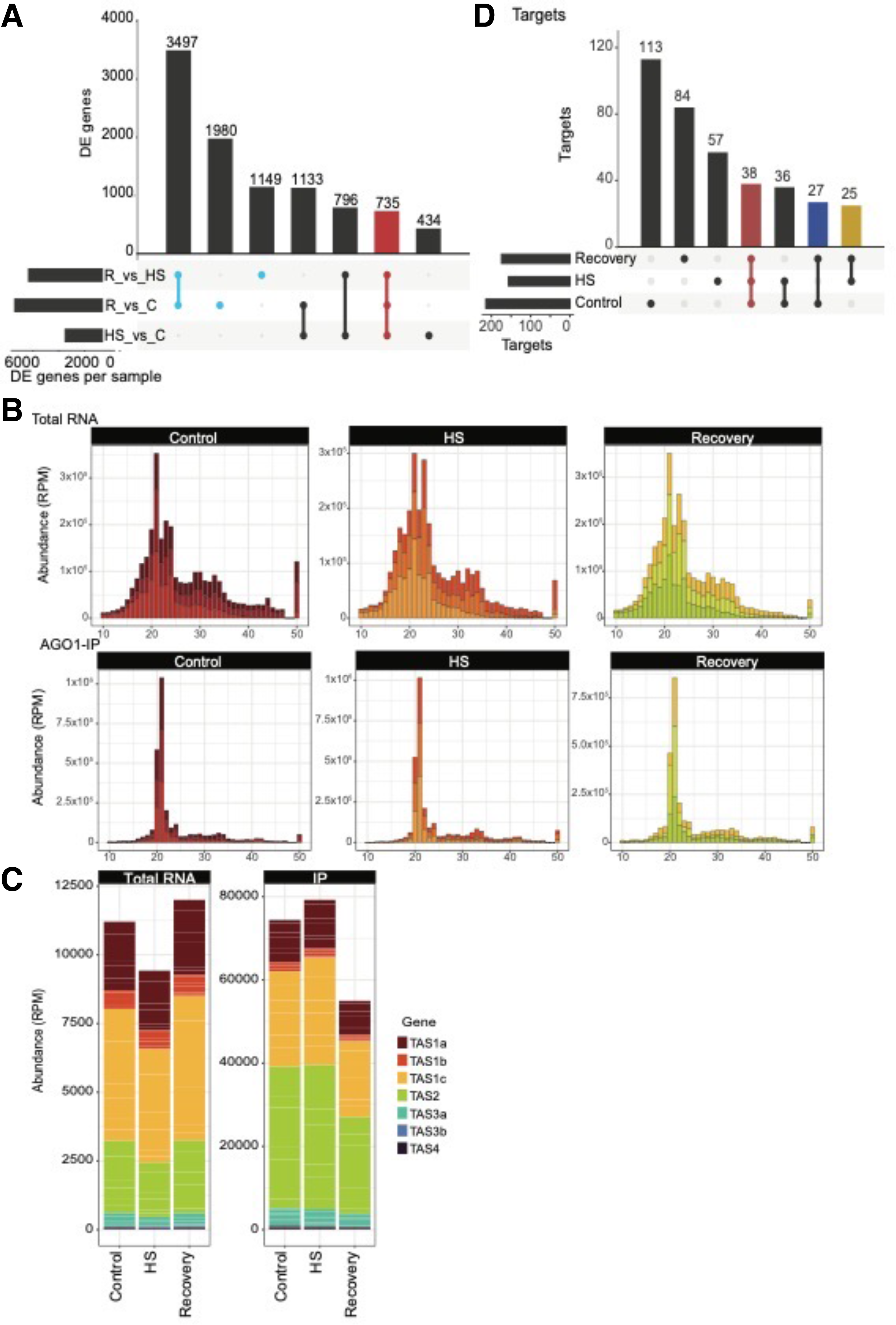
Differentially expressed coding genes and sRNA accumulation during HS and recovery. (A) Upset plot representing statistically significant differentially expressed (DE) genes between all three comparisons. (B) Size distribution of sRNAs mapping to the Arabidopsis genome (TAIR version 10). The abundance of each size class was normalized to reads per million (RPM) for each library. The x-axis indicates the sRNA size and the y-axis indicates its abundance in RPM-mapped reads. Shown are data from three independent biological replicates, for total RNA (upper panel) and AGO1-IP (lower panel). (C) Abundance of each TAS gene in Total RNA (left panel) and AGO1-IP (right panel). The x-axis represents the sample type, and the y-axis represents the abundance of sRNAs mapping each TAS gene. The three biological replicates for each treatment are plotted and stacked together. Each TAS gene is represented with a different color. (D) Upset plot representing the miRNA targets identified by nanoPARE sequencing in each of the treatments. For each upset plot, the bottom left shows the number of differentially expressed genes in each comparison as a horizontal histogram, the bottom right shows the intersection matrix and the upper right shows the size of each combination as a vertical histogram. The red line marks differentially expressed genes common to all three comparisons, the blue lines mark differentially expressed genes during the recovery process.

**Fig. S6.**
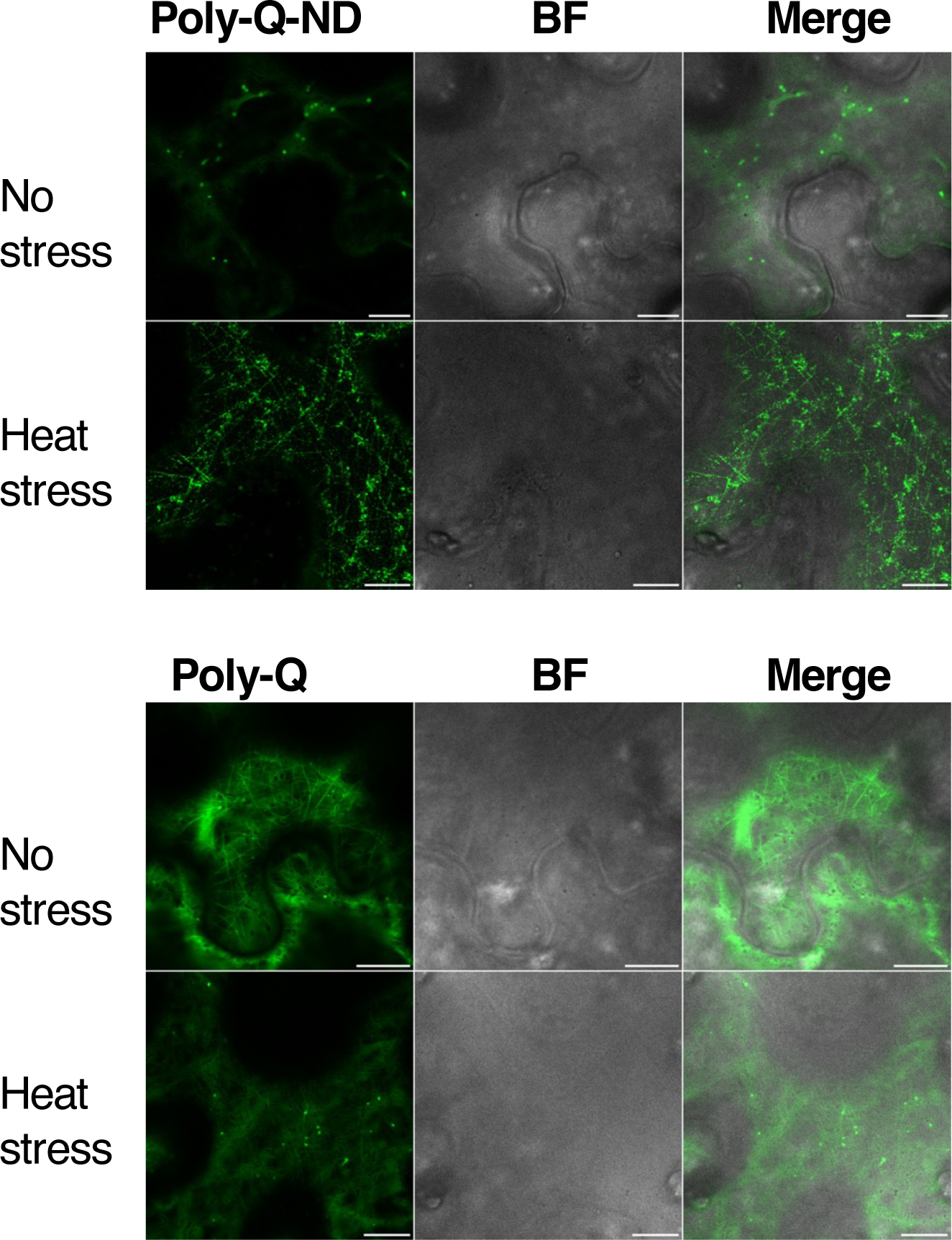
The Poly-Q domain alone undergoes LLPS in plant cells under HS. Representative CLSM imaging of *N. benthamiana* leaf epidermal cells transiently expressing GFP-Poly-Q-ND and GFP-Poly-Q proteins in the absence or presence of HS (30 minutes at 37°C). Bar = 10 µm. Objective 40X, oil immersion. BF, bright field.

**Fig. S7.**
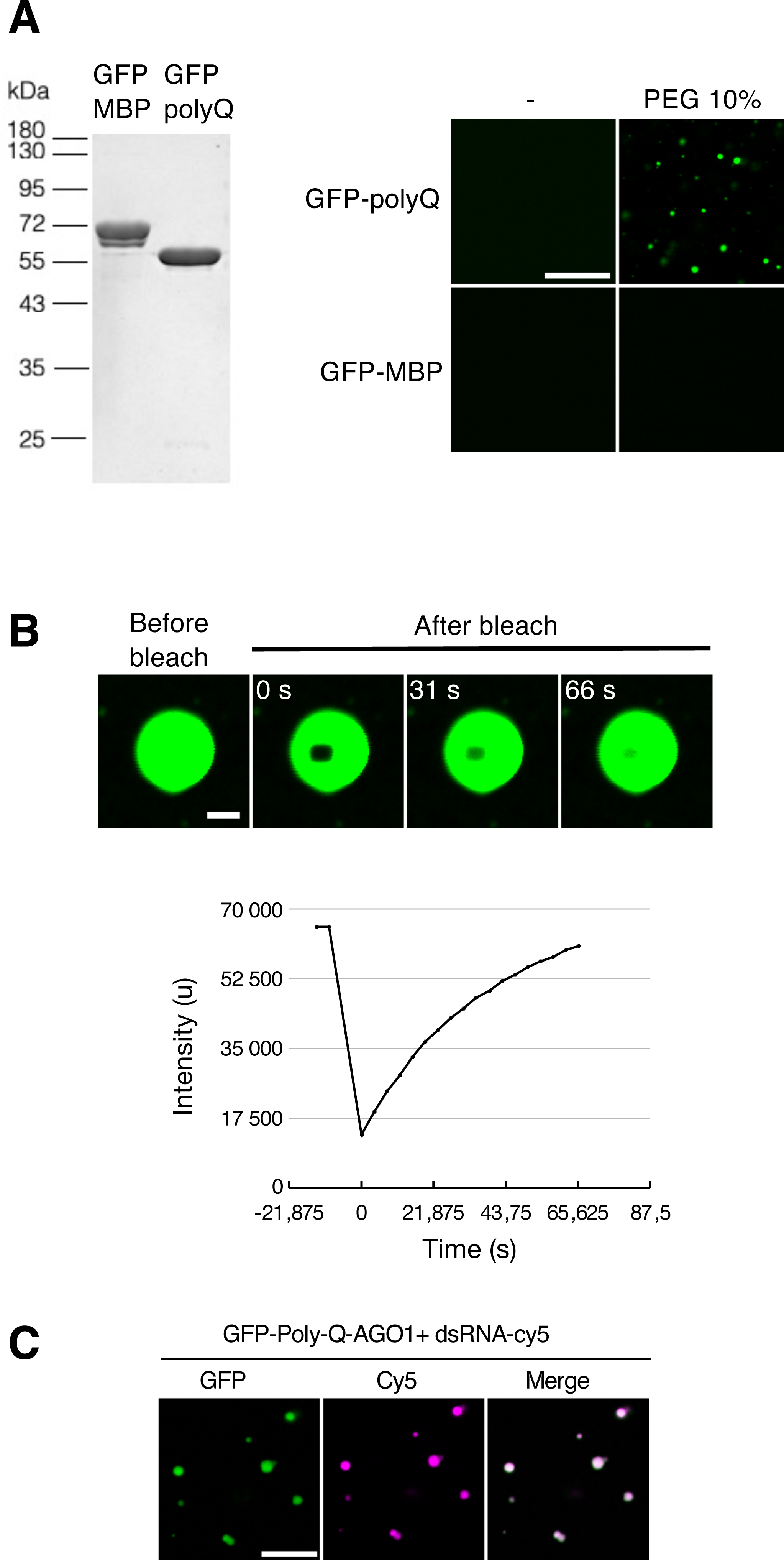
The Poly-Q domain alone undergoes LLPS and colocalize with RNA in vitro. **(A)** AGO1 Poly-Q domain, but not maltose binding protein fused to GFP form droplets *in vitro*.. Left panel: SDS-PAGE analysis of the recombinant protein GFP-polyQ AGO1 protein and GFP-MBP used in *in vitro* LLPS assays and EMSA assays. 2,5ug per lane of 12% Tris-glycine SDS-PAGE. Right panel: *In vitro* analysis of droplet formation by recombinant GFP-polyQ AGO1 protein and control protein GFP-MBP protein at 200mM NaCl concentration in Hepes 25mM buffer pH7.5, PEG 8000 10%. Protein concentration 4μM Scale bars, 20μm. Objective 20X. **(B)** FRAP of GFP-Poly-Q AGO1 droplets. Time 0 is set at the time of the photobleaching pulse. Data are representative of four independents experiments. Scale bar, 2μm. At the bottom, recovery data corresponding with images displayed in the upper part. **(C)** Colocalization of GFP-Poly-Q AGO1 protein with Cy5 labelled-dsRNAs upon droplet formation in Hepes 25mM buffer pH7.5, NaCl 200mM, PEG8000 10%. Protein and RNAs are at 4μM. Scale bars, 10μm. Objective 40X.

**Fig. S8.**
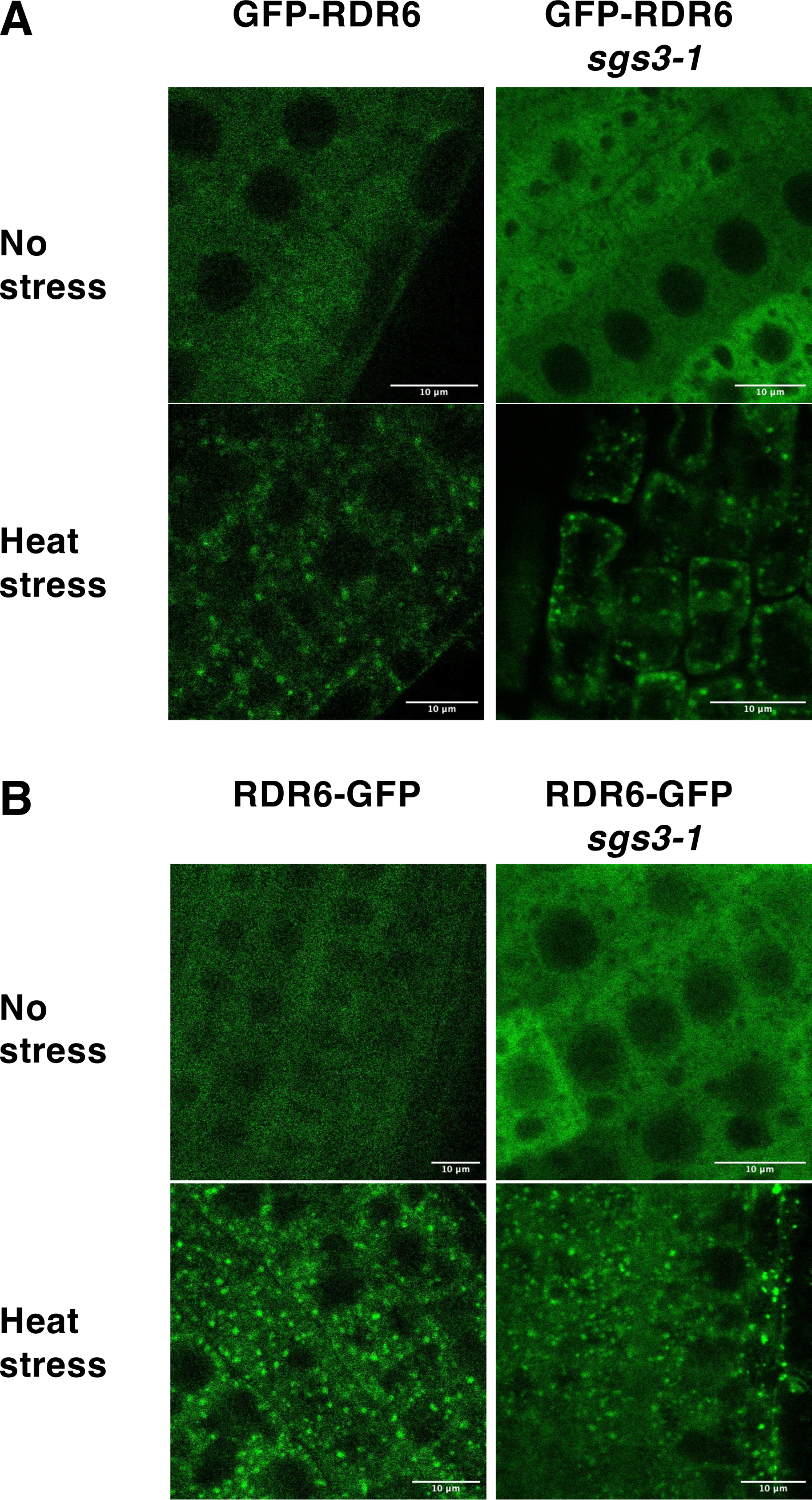
HS promotes GFP-RDR6 foci formation even in the absence of *SGS3*. **(A)** Representative CLSM imaging of 5 day old Arabidopsis root tip cells of the p35S:GFP-RDR6 in Col-0 or *sgs3-1* transgenic line subjected to 60 min HS at 37°C. Bar = 10 µm. Objective 60X, water immersion. **(B)** Representative CLSM imaging of 5 day old Arabidopsis root tip cells of the p35S:RDR6-GFP in Col-0 or *sgs3-1* transgenic line subjected to 60 min HS at 37°C. Bar = 10 µm. Objective 60X, water immersion.

